# Perturbation-Specific Transcriptional Mapping for unbiased target elucidation of antibiotics

**DOI:** 10.1101/2024.04.25.590978

**Authors:** Keith P Romano, Josephine Bagnall, Thulasi Warrier, Jaryd Sullivan, Kristina Ferrara, Marek Orzechowski, Phuong H Nguyen, Kyra Raines, Jonathan Livny, Noam Shoresh, Deborah T Hung

## Abstract

The rising prevalence of antibiotic resistance threatens human health. While more sophisticated strategies for antibiotic discovery are being developed, target elucidation of new chemical entities remains challenging. In the post-genomic era, expression profiling can play an important role in mechanism-of-action (MOA) prediction by reporting on the cellular response to perturbation. However, the broad application of transcriptomics has yet to fulfill its promise of transforming target elucidation due to challenges in identifying the most relevant, direct responses to target inhibition. We developed an unbiased strategy for MOA prediction, called Perturbation-Specific Transcriptional Mapping (PerSpecTM), in which large-throughput expression profiling of wildtype or hypomorphic mutants, depleted for essential targets, enables a computational strategy to address this challenge. We applied PerSpecTM to perform reference-based MOA prediction based on the principle that similar perturbations, whether chemical or genetic, will elicit similar transcriptional responses. Using this approach, we elucidated the MOAs of three new molecules with activity against *Pseudomonas aeruginosa* by comparing their expression profiles to those of a reference set of antimicrobial compounds with known MOAs. We also show that transcriptional responses to small molecule inhibition resemble those resulting from genetic depletion of essential targets by CRISPRi by PerSpecTM, demonstrating proof-of-concept that correlations between expression profiles of small molecule and genetic perturbations can facilitate MOA prediction when no chemical entities exist to serve as a reference. Empowered by PerSpecTM, this work lays the foundation for an unbiased, readily scalable, systematic reference-based strategy for MOA elucidation that could transform antibiotic discovery efforts.

**Significance Statement:** New antibiotics are critically needed in the face of increasing antibiotic resistance. However, mechanism-of-action (MOA) elucidation remains challenging and imposes a major bottleneck in antibiotic discovery and development. Building on the principle that molecules with similar MOAs elicit similar transcriptional responses, we have developed a highly scalable strategy for MOA prediction in the important bacterial pathogen *Pseudomonas aeruginosa* based on correlations between the expression profiles of new molecules and known perturbations, either small molecule inhibition by known antibiotics or transcriptional repression of essential targets by CRISPRi. By rapidly assigning MOAs to three new molecules with anti-pseudomonal activity, we provide proof-of-concept for a rapid, comprehensive, systematic, reference-based approach to MOA prediction with the potential to transform antibiotic discovery efforts.

## Introduction

*Pseudomonas aeruginosa* and other Gram-negative bacteria pose a major threat to human health due to the growing prevalence of antibiotic resistance in healthcare settings. Surging multi-drug-resistant bacterial infections threaten to outpace our ability to develop novel antibiotics [1]. Antibiotic discovery campaigns have failed to deliver new antibiotic classes, including extensive efforts in the post-genomic era. As recently reviewed, new strategies are being developed to address the looming antibiotic shortage, including novel approaches to mine the natural products space [5–7], to apply artificial intelligence [8], and to leverage genetic engineering of strains in a chemical systems biology approach [9]. Most of these novel approaches differ from standard *in vitro*, target-based strategies in which the target is known by design, as they prioritize the identification of new molecules with whole-cell antibacterial activity. While accelerating the discovery of new chemical entities, these modern approaches highlight the downstream challenge of mechanism-of-action (MOA) determination and the need for better, more systematic methods to tackle this problem to enable candidate development. Currently, extensive biochemical and genetic efforts are required for target and MOA elucidation, which impose a major bottleneck in antibiotic discovery and development pipelines [10]. Indeed, early MOA assignment for antibacterial small molecule candidates would provide mechanistic and biological insights to guide their prioritization, medicinal chemistry optimization, and eventual hit-to-lead transition.

Transcriptional profiling has long been used to explore the MOA of antimicrobial agents, first described in yeast using microarrays to delineate the MOA of the inhibitor dyclonine [2], and later performed in a wide array of bacteria, including *Bacillus subtilis*, *Mycobacterium tuberculosis*, *Staphylococcus aureus*, *Escherichia coli and P. aeruginosa* [3–9]. Work in *B. subtilis* generated the first antimicrobial reference set based on gene expression using microarrays to characterize the transcriptional responses to 37 antibiotics [3]. More recently, work in *E. coli* also included a reference set of 37 antibiotics, from which the authors chose gene sets based on differential gene expression analysis of compound treatment relative to negative control, and used correlation coefficients to predict MOAs [6]. Then, Espinoza et al. furthered this work by developing a machine learning approach for transcriptomics-based MOA prediction in *E. coli*, which involved a reference set of 41 antibiotics and a feature selection algorithm called *Clairvoyance* [7].

Despite these advances, transcriptomics-based MOA prediction remains limited by the challenge of extracting meaningful signals that distinguish the relevant gene responses to target inhibition from less specific transcriptional changes [10, 11]. Ideally, transcriptomics-based MOA prediction should be rooted in gene expression responses specific to the given chemical or genetic perturbation. However, during perturbation, bacteria also change the expression of a plethora of background genes, which may be far downstream from the perturbed pathway or unrelated altogether, which obfuscate the perturbation-specific signals. For example, nonspecific gene responses, relating to cellular stress or impending cell death, can be shared across antimicrobials compounds with disparate MOAs. Meanwhile, differences in measured expression programs can also be related to growth phase differences between treated and untreated cells as they grow or divide at different rates, rather than a direct effect of antimicrobial activity. To manage these effects, treatments with sublethal concentrations of compound are often used to minimize the growth arrest or death of treated populations and match growth rates of treated and untreated samples; however, these low treatment concentrations often yield weak transcriptional responses, thereby obscuring informative, relevant changes that occur with compound exposure [12–18].

To address this challenge, we developed a systematic method called Perturbation-Specific Transcriptional Mapping (PerSpecTM) that combines experimental and analytic methods that leverage its large-throughput scale to accentuate changes in gene expression that are specific to and a direct consequence of target inhibition. By using a diverse set of perturbed transcriptomes for normalization, an approach adapted from large-scale gene profiling in human cancer cell lines [22], PerSpecTM emphasizes perturbation-specific transcriptional signatures while deemphasizing common, nonspecific responses. It then assigns target and MOA in a reference-based manner by correlating the pattern of changes with the transcriptional responses of a reference set, thus avoiding biases that might occur by assigning MOA based on the response of specific, predetermined genes [16, 19].

We applied PerSpecTM to the important bacterial pathogen *P. aeruginosa*, building two complementary reference sets: (1) an antimicrobial reference set comprising 37 antimicrobial compounds; and (2) a genetic depletion reference set comprising 14 strains, each exhibiting tunable transcriptional repression of a different essential gene target using CRISPRi. Querying against the antimicrobial reference set, we rapidly elucidated the targets and MOAs of three new molecules with anti-pseudomonal activity, while demonstrating the value of using genetically engineered, hypersensitized strains depleted for essential targets to yield more robust transcriptional responses to perturbation. Because of the limited number of chemical entities with known unique targets, and thereby MOAs, that can serve as an antimicrobial reference, we applied PerSpecTM to the transcriptional repression reference set to demonstrate proof-of-concept that genetically repressing target expression can phenocopy its chemical inhibition, thus affording target and MOA prediction when no chemical entities exist to serve as a reference. Empowered by PerSpecTM and mutant strains depleted in essential targets, this work enables an unbiased and highly scalable reference-based strategy for target and MOA elucidation to assist antimicrobial discovery efforts in the face of the looming shortage of antibiotics in healthcare systems worldwide.

## Results

### Developing Perturbation-Specific Transcriptional Mapping (PerSpecTM)

We set out to generate in large-throughput, an antimicrobial reference set of chemical treatments of the *P. aeruginosa* laboratory strain UCBPP-PA14 (abbreviated hereafter as PA14) and related mutant strains. To enable such scale, we worked to determine the optimal assay conditions in 384-well format and analysis strategy in a manner that would not require tailoring conditions for each individual compound.

All expression profiling experiments were performed in 384-well format with 60μL per well of bacterial culture at an inoculum of ∼1 × 10^8^ colony-forming unit (CFU) per milliliter (mL), used to ensure that sufficient cells were available to guarantee the generation of a high-quality RNAseq library. Each compound was dosed in triplicate at 3-fold the minimum-inhibitory concentrations (MICs), with the MIC determined by conventional broth microdilution at a much lower inoculum of ∼5 × 10^5^ CFU/mL [12]. Plates were arrayed with a diverse set of active compounds across multiple MOA classes, as well as DMSO and water controls. Resulting gene expression data for each gene of each sample were standardized (transformed to z-scores) with respect to all samples in the same batch, where each batch is defined as samples on the same 384-well plate sharing the same strain and time point. The z-score provides a measure of how unusually high or low the expression of a given gene is after a specific chemical (or genetic) perturbation relative to its expression across the entire population of samples within a batch. Provided the batch consists of a diverse set of active antimicrobial agents (or genetic perturbations), z-scores highlight transcriptional changes that are unique to a given perturbation, and deemphasize common gene responses across the population, such as those relating nonspecifically to growth, cellular stress, or impending cell death. Thus, the relevant signals are accentuated in an unbiased manner, without needing to subset genes based on computational approaches or *a priori* knowledge of expected gene responses. The ability to calculate z-scores across a variety of treatments, rather than just assessing differential expression between a particular sample and negative control samples, was made possible by the large-throughput scale of PerSpecTM. To then assess overall strength of the transcriptional signals, differential gene expression was orthogonally assessed using DESeq2 [68]. A transcriptionally weak signal was defined as the differential expression between compound treatment and vehicle treatment wherein no gene had an adjusted p-value that was more significant than the minimum adjusted p-value found when comparing PA14 in water versus DMSO control (i.e. p_adj_ < 10^−20^). Although weak signals may reflect meaningful biological changes, we wanted to only include robust signals in the antimicrobial reference set to facilitate reliable target predictions. Similarities between perturbations were then calculated using the Pearson correlation coefficients between transcriptional signatures of two conditions, where each gene’s expression is represented as a z-score.

We first set out to determine the optimal incubation time and compound concentration yielding robust transcriptional responses across many different compound treatments, without individual compound optimization. We profiled a set of 21 antimicrobial compounds at 30, 60, 90, or 120 minutes, depicted in bold in **Table S1**, which collectively targeted four distinct cellular mechanisms: inhibitors of DNA synthesis, protein synthesis, cell wall (defined as the structural elements of the cell envelope, excluding the cell membranes), and membrane integrity. After calculating z-scores for each gene and determining the Pearson distances (√(1 − *Pearson Correlation*)) between adjacent timepoints over the 120-minute time trial (**SI Appendix, Fig. S1a**), we found that the transcriptional responses to treatment by inhibitors of DNA synthesis, protein synthesis and membrane integrity converged on transcriptional endpoints by 90 minutes within each MOA. The transcriptional responses to treatment with cell wall synthesis inhibitors were more heterogenous as reflected in their larger Pearson distances, which never fully converged by 120 minutes.

We wondered if the lack of transcriptional convergence of cell wall inhibitors by 120 minutes related to the inoculum effect, a phenomenon observed in β-lactams causing loss of activity at high bacterial densities [13]. We measured the MICs of compounds at higher inoculum (i.e. 1×10^8^ CFU/mL used in RNAseq experiments) compared to the lower convention of ∼5 × 10^5^ CFU/mL [12]. **Table S1** denotes the antimicrobial compounds exhibiting strong inoculum effects (≥ 4-fold shift in MIC at the high bacterial density) with an asterisk, which comprise all of the cell wall synthesis inhibitors, except imipenem, due to upregulation of the beta lactamase AmpC at high bacterial density [14]. Interestingly, the Pearson distances for imipenem across the time trial also converge by 90 minutes, comparable to inhibitors of protein synthesis, DNA synthesis and membrane integrity (**SI Appendix, Fig. S1a**). Taken together, these data suggested that for antimicrobial compounds which are active at the experimental inoculum, an incubation time of 90 minutes yielded a stable transcriptional endpoint.

After establishing an optimal 90-minute incubation time, we sought to determine the optimal antimicrobial concentration that elicits a strong transcriptional response. Here we used an expanded set of 39 antimicrobial compounds, shown in **Table S1**, spanning the same four MOAs but through inhibition of more diverse targets. We added several compounds with known anti-pseudomonal activity, but not yet approved for human use. For example, we included cell wall-related compounds, such as inhibitors of LpxC, LptD, BamA and MreB that disrupt lipopolysaccharide (LPS) biosynthesis, LPS transport, outer membrane protein assembly, and cell shape, respectively. We also included the non-pseudomonal antimicrobial compound nitrofurantoin, which works through a complex mechanism involving reactive oxygen species that interfere nonspecifically with DNA synthesis, as well as protein and cell wall metabolism [15]. We also added: (1) the antibacterial agent fosmidomycin, which inhibits Dxr to block isoprenoid synthesis [16]; (2) the Hsp90 inhibitor radicicol, an antifungal agent that likely interferes with gyrase-related proteins with a conserved Bergerat fold [17–19]; (3) the anti-staphylococcal agent fusidic acid, an inhibitor of the elongation factor FusA1 that blocks translation [20]; (4) the antibiotic rifampicin, which blocks transcription through the inhibition of the DNA-dependent RNA polymerase RpoB [21]; (5) the fatty acid synthesis inhibitor thiolactomycin; and (6) the membrane pore former LL-37 [22]. Lastly, we included several anti-cancer agents, such as cisplatin, doxorubicin and hydroxyurea, which exhibit antimicrobial activity by inhibiting DNA synthesis through DNA alkylation, deoxyribonucleic acid depletion and DNA intercalation, respectively [23–29].

Compounds were tested at 0.5-fold, 2-fold and 4-fold their adjusted MICs as determined in broth microdilution experiments at high inoculum, thereby accounting for any inoculum effects. In some cases, the highest possible concentrations were limited by solubility. These data confirmed that all compounds dosed above their MICs demonstrated robust transcriptional responses (**SI Appendix, Fig. S1b**). In contrast, antimicrobials that were inactive (non-lethal) at the applied concentrations yielded variable transcriptional responses. In some cases, such as for DNA synthesis inhibitors and membrane disruptors, robust gene expression signals were generated even under subinhibitory concentrations, suggesting that bacteria are primed to detect non-lethal perturbations of certain cellular processes. In the case of some of the β−lactam antibiotics, such as piperacillin, ceftazidime, and aztreonam, robust transcriptional responses could not be elicited due to compound insolubility at concentrations necessary to overcome the inoculum effect (**SI Appendix, Fig. S1c**). In fact, because aztreonam treatment yielded a transcriptionally weak response across all treatment conditions, it was removed from the reference set moving forward. Taken together, these results demonstrate that antimicrobial activity at the high experimental inoculum (i.e. 1 × 10^8^ CFU/mL) was necessary to ensure a robust transcriptional response to all antibiotics in this large-throughput assay format, even while some compounds are still able to generate robust signals despite sublethal dosing. We thus settled on the general use of 2-4X the MIC measured at a high inoculum for compound treatments.

### Transcriptional responses to antimicrobial compounds with similar MOAs are correlated

As expected, our large-throughput gene expression dataset confirmed that lethal antimicrobial treatments elicited transcriptional responses that correlated by MOA. The Pearson correlations of z-scores of pairwise combinations of 90-minute treatments, at the highest dose for each compound, are shown as a similarity matrix in **Fig. 1a**. Compounds with limited anti-pseudomonal activity, comprising doxorubicin, radicicol, hydroxyurea, piperacillin, and rifampicin, had positive correlation values with vehicle treatment. Of the remaining antimicrobial compounds, Pearson correlations were strongest between compounds sharing the same cellular mechanism. Not only did compounds correlate by MOAs, but stronger correlations were observed within antibiotic subclasses. For example, based on the strength of their correlations, protein synthesis inhibitors grouped into macrolides (clarithromycin and erythromycin), tetracyclines (tetracycline, minocycline and doxycycline) and aminoglycosides (amikacin and gentamicin). Likewise, fluoroquinolones (ciprofloxacin and levofloxacin) formed a clear subgroup within the larger class of DNA synthesis inhibitors. Finally, the transcriptional responses to LptD inhibitors (POL-7001 and POL-7080) and LpxC inhibitors (PF-5081090 and PF-04753299), which block LPS synthesis and transport to the cell surface, respectively, tightly correlated within the group of membrane integrity inhibitors. Meanwhile, despite the resolution obtained for some subclasses, inhibitors of membrane integrity and cell wall synthesis elicited transcriptional responses that cross-correlated, pointing to a shared cellular response to defects in membrane disruption and cell wall synthesis.

**Fig. 1.**
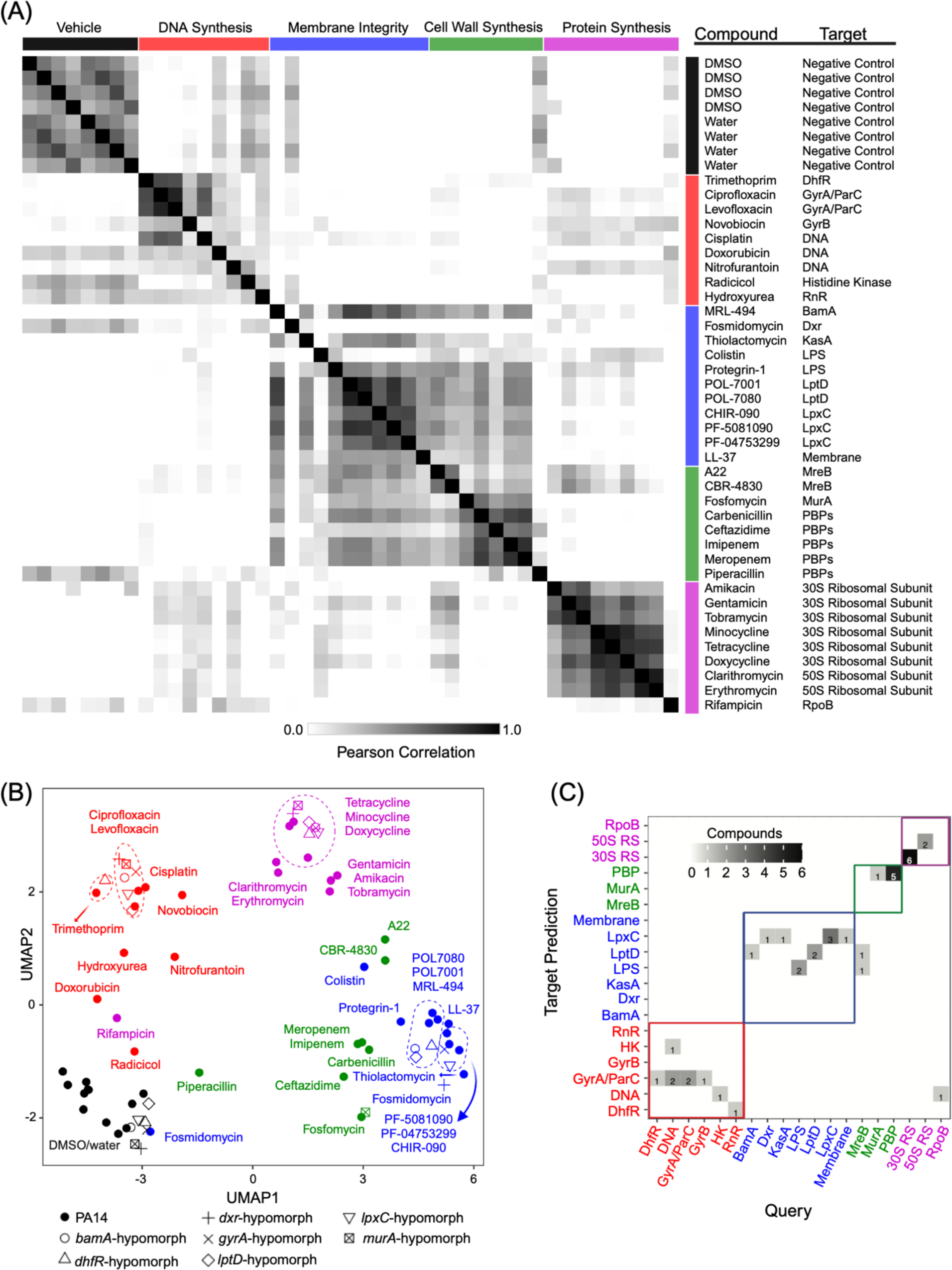
Correlating antimicrobial gene expression profiles of the antimicrobial reference set. (A) Heat map of Pearson correlations of the expression profiles of PA14 treated with all 37 compounds in the antimicrobial reference set at the highest concentration at 90 minutes. Rows and columns are organized and color-coded by MOA. All correlation values below zero were set to zero for visualization purposes. (B) UMAP visualization of all compound treatments (at the highest dose) of wildtype PA14 and seven engineered hypomorphs at 90 minutes. Colors correspond to antibiotic MOA classes: DNA synthesis (red), membrane integrity (blue), cell wall synthesis (green), and protein synthesis (magenta). Closed circles indicate wildtype PA14 treatments, whereas all other markers indicate treatment of a hypomorphic strain, as indicated. Antimicrobial compounds with robust activity at the experimental inoculum group by MOA, agnostic of the genetic strain. Protein synthesis inhibitors form subgroups that separate tetracyclines, macrolides and aminoglycosides. Except for novobiocin, compounds that are not active against PA14, such as fosmidomycin, piperacillin, rifampicin, doxorubicin and hydroxyurea, appear close to DMSO vehicle. Treatments that were deemed transcriptionally weak based on DESeq2 analysis were not included in the visualization. (C) Confusion matrix showing the number of compounds in each query target category that were predicted to have indicated targets in the leave-one-out cross validation (LOOCV), using the method of calculating the target correlation prediction score (*r*_*predict*_). Compounds with correctly predicted target fall along the diagonal, while those with correctly predicted MOA fall within the colored boxes. Note that query target categories that only contain one compound (the column sums to one) could not be predicted correctly, due to the leave-one-out approach.

Lastly, to verify that the transcriptional responses to compound treatment corresponded with known biology, we determined the most upregulated and downregulated genes that emerged from calculating z-scores in response to three of the most well-studied antibiotic classes: aminoglycosides, fluoroquinolones and polymyxins. **Table S2** shows the top 25 upregulated and downregulated genes with known annotations in PA14 in response to treatment with aminoglycosides, fluoroquinolone and polymyxins. Indeed, previously reported gene expression signatures were readily apparent for each antibiotic class, providing a sanity check that the PerSpecTM approach indeed highlights changes in gene expression directly related to each specific chemical perturbation.

### Transcriptional responses to treatment of hypersusceptible strains correlate with wildtype treatments

The requirement that compounds have potent wildtype activity to elicit a reliable, robust, informative transcriptional response poses a barrier to the early integration of transcriptomics-based MOA prediction for newly discovered compounds which might not yet have highly potent antimicrobial activity. To address this concern, we sought to determine whether mutant strains that are hypersusceptible to compounds could be used in place of wildtype strains to obtain robust transcriptional signals for compounds with limited wildtype activity. While an efflux pump mutant could serve as a hypersusceptible strain to some compounds, not all compounds have increased sensitivity in such a strain. We thus selected an alternative strategy involving the engineering of hypomorphic mutants that are hypersensitized to inhibitors by depleting their cognate gene products[30].

To test this concept, we sought to create hypomorphs and compare their transcriptional responses to antimicrobial compounds with those of wildtype bacteria. We genetically engineered seven hypomorphic strains using a regulated promoter replacement strategy to deplete essential gene targets corresponding to a set of known antimicrobial agents (**SI Appendix**, **Additional Methods** for details of strain construction). As summarized in **Table S3**, the panel of hypomorphic strains compromised gene target depletions of the LPS transporter protein LptD, the isoprenoid synthesis protein Dxr, the peptidoglycan synthesis protein MurA, DNA gyrase GyrA, the LPS biosynthesis enzyme LpxC, the dihydrofolate reductase DhfR involved in folate biosynthesis, and the outer membrane protein assembly protein BamA. The growth phenotypes were unchanged for most engineered hypomorphs, but reduced in the case of *gyrA*, *dxr* and *lptD* hypomorphs (**SI Appendix**, **Fig. S2**). We then measured the MICs of compounds targeting each depleted gene product: POL7080 against LptD [31], fosmidomycin against Dxr [16], fosfomycin against MurA [32], ciprofloxacin against GyrA [33], PF-04753299 against LpxC [34], trimethoprim against DhfR [35], and MRL-494 against BamA [36]. The *lptD*, *murA*, and *bamA* hypomorphs were 15–20-fold more susceptible to their cognate inhibitors, whereas the *dxr*, *gyrA*, *lpxC* and *dhfR* hypomorphs showed no significant shifts in MIC relative to wildtype (**Table S3**). Except for the *lptD*-hypomorph against doxycycline, the hypomorphs did not show appreciable hypersensitivity to inhibitors whose MOA were unrelated to the specific target depleted in each strain.

We measured the gene expression profile of each hypomorph treated with vehicle, one on-target compound (*i.e.,* compound whose activity is due to inhibition of the corresponding target) and two off-target compounds (*i.e.,* compound whose activity is not related to the corresponding target). Hypomorphic strains were incubated with compounds at 2-fold their MICs (determined at high inoculum against each hypomorph), or vehicle. Z-scores were calculated across each hypomorph, rather than across all hypomorphs, to dampen background transcriptional changes related to strain-specific repression in gene expression. **Fig. 1b** shows a combined uniform manifold approximation and projection (UMAP) showing wildtype and all mutants with their corresponding treatments. The changes in gene expression of hypomorphs to chemical perturbation correlated with wildtype treatment responses. For example, treatment of the g*yrA*-hypomorph with its on-target inhibitor, ciprofloxacin, grouped with PA14 treatment with ciprofloxacin, as was the case for *lpxC*-hypomorph treated with PF-04753299, *bamA*-hypomorph treated with MRL-494, *dhfR*-hypomorph treated with trimethoprim, *murA*-hypomorph treated with fosfomycin, and *lptD*-hypomorph treated with POL7080. Meanwhile, grouping of mutant and wildtype responses was agnostic of whether a compound’s molecular target related to the underlying genetic alteration in a strain, as exemplified by the treatment response of the *gyrA*-hypomorph to doxycycline, a protein synthesis inhibitor unrelated to GyrA, grouping together with PA14 in response to doxycycline.

The correlation between (separately standardized) responses of hypersusceptible mutants and wildtype enabled the characterization of expression responses of compounds without wildtype activity. Indeed, the transcriptional response of the *dxr-*hypomorph to fosmidomycin provides proof-of-concept for the utility of using genetically depleted strains to facilitate MOA prediction. Fosmidomycin inhibits Dxr, an essential reductoisomerase in the 2-C-methyl-D-erythritol 4-phosphate (MEP) pathway of isoprenoid precursor biosynthesis [37]. While fosmidomycin exhibited an MIC of 12.5μM at low inoculum, it showed no activity against PA14 at the high inoculum used for expression profiling experiments. In contrast, fosmidomycin caused a significant growth delay on the *dxr*-hypomorph at high inoculum, with an IC_50_ of about 25μM (**SI Appendix**, **Fig. S3**). While the expression response of wildtype PA14 to fosmidomycin treatment grouped near vehicle controls, the response of the *dxr*-hypomorph grouped with inhibitors of membrane integrity, accurately reflecting the cellular mechanism of fosmidomycin **(Fig. 1b).**

### MOA and target prediction using the antimicrobial reference set

The antimicrobial reference set was curated to include robust transcriptional profiles of all aforementioned treatments across various time points, doses and hypomorphic strains. Treatment conditions eliciting weak transcriptional signals were excluded, which fully excluded aztreonam from the reference set, as described earlier. In addition, any treatment condition having a z-score transcriptional profile that correlated more highly to negative control samples than to any other condition in the data set, including other treatment conditions of the same compound, were also removed. This resulted in the exclusion of fusidic acid from the reference set. In total, 37 compounds were retained in the antimicrobial reference set.

We next set out to evaluate the ability of PerSpecTM to perform reference-based target and MOA predictions for novel chemical entities using an antimicrobial reference set, with target referring to the biomolecule engaged by a given compound, and MOA referring to the cellular function disrupted by target engagement. We hypothesized that the predicted target for a query compound could be determined by the highest overall Pearson correlation between the query compound and all other treatments within the reference set, similar to methods described previously [6]. In practice, to arrive at a single target prediction for a given compound, in cases where there are multiple treatment conditions (*e.g.,* multiple doses or time points), we first determined the maximal Pearson correlation (*r*_*max*_) between each query treatment and every individual treatment within each reference target category. We removed query treatments whose highest *r*_*max*_ derived from negative control samples. Then, we averaged the *r*_*max*_ values from all remaining query treatments for a given compound to define a target correlation score, *r̅*_*max*_. In the case where there is only one treatment condition for a query compound, *r̅*_*max*_ = *r*_*max*._. The final target prediction was assigned based on the highest *r̅*_*max*_ across all target categories in the reference set, which we termed the target correlation prediction score, *r*_*predict*_. In the case where there is only one treatment condition for a query compound, *r*_*predict*_ is the equivalent to the highest overall Pearson correlation between the query compound and all reference treatments. The predicted MOA of the query compound was then assigned as the known cellular pathway associated with the predicted target.

To evaluate the performance of PerSpecTM, we performed a leave-one-out cross validation (LOOCV) (**Fig. 1c**), wherein the expression profiles for each individual antimicrobial compound in the reference set were queried against the expression profiles of all other compounds in the reference set. The correlation analysis was carried out as described above, correctly assigning MOAs for 34 of the 37 antimicrobial compounds (92%), and correctly assigning targets for 22 of the compounds (60%). Of note, 10 compounds (27%) were the only representatives of their respective target categories, and thus LOOCV had no chance of correct target prediction. Removing those ten singleton compounds from the query, the correct MOAs were assigned for 25 out of 27 compounds (94%), and correct targets were predicted for 22 of the compounds (81%).

Next, to assess confidence in each MOA and target prediction, we obtained positive predictive values (PPVs) for both MOA and target assignments, respectively. The PPV is the percent of compounds in the LOOCV whose prediction was correct for *r*_*predict*_ values above a given threshold. Of note, the distributions of *r*_*predict*_are different for different MOAs, and thus may merit different levels of confidence depending on the predicted MOA (**SI Appendix, Fig. S4**). However, due to the limited number of compounds per category, the PPVs were calculated across the entire data set for both MOA and target. For each target and MOA prediction in the subsequent sections, we report the respective PPVs that are associated with the final prediction score, *r*_*predict*_.

### External validation of the antimicrobial reference set

To test PerSpecTM’s performance on external data, we queried the largest publicly available gene expression dataset in response to antibiotic and antiseptic treatments in *P. aeruginosa* [9]. Notably, the experimental design of the query dataset was highly divergent from the methods employed in our study. Gene expression experiments comprised sublethal treatments in duplicate with 14 antimicrobial compounds, including one fluoroquinolone (ciprofloxacin), three aminoglycosides (gentamicin, neomycin, and tobramycin), three β−lactams (ampicillin, carbenicillin and cefoperazone), one membrane disruptor (polymyxin B), one cell wall synthesis inhibitor (aztreonam) and five antiseptics (benzalkonium chloride, bleach, peroxide, povidone-iodine and silver nitrate). The size and chemical diversity of the query dataset allowed us standardize gene expression with respect to the entire query dataset. Ampicillin, aztreonam, carbenicillin, cefoperazone, povidone-iodine, silver nitrate, benzylalkonium chloride and tobramycin were removed from the analysis, due to either weak transcriptional signals, or responses that correlated most to negative controls in the reference set. **Table S4** summarizes the MOA and target predictions of the 6 remaining external chemical treatments included in the analysis. Although neomycin had a transcriptionally weak response, it was nevertheless included in the analysis because its *r*_*predict*_ value was within the range of those in the LOOCV. Of the six external compounds included in the analysis, two were also in our antimicrobial reference set (ciprofloxacin and gentamicin), two had MOAs represented in our reference set even though the exact compounds were not present in the reference set (neomycin and polymyxin B), leaving two (bleach and peroxide) as completely unique to the external dataset.

Notably, the MOA was predicted correctly for four of the six external compounds in the analysis (**Fig. 2a**). The MOA and target were correctly predicted for the two external compounds that were also in our reference set: gentamycin was correctly assigned as a protein synthesis inhibitor with the 30S ribosomal subunit as the predicted target; and ciprofloxacin was correctly assigned as a DNA synthesis inhibitor with GyrA as the predicted target. This indicates that the standardization method was able to compensate for the technical differences between the datasets. Correct MOAs were also assigned for the two external compounds not in the reference set, but whose targets and MOAs were represented by other compounds in the reference set. Neomycin was correctly predicted to be a protein synthesis inhibitor of the 30s ribosomal subunit. While no confident predictions could be made for the target or MOA of polymyxin B due to the low *r*_*predict*_ value, the correct MOA was implicated as membrane integrity disruption. Polymyxin’s highest target correlation score, *r̅*_*max*_, was with LptD, while the second highest *r̅*_*max*_ corresponded to LPS, its known target (**Fig. 2a**). Interestingly, of the antiseptics, peroxide was assigned as a DNA synthesis inhibitor with GyrA and DNA as the top two predicted targets, similar to fluoroquinolones and cisplatin. Meanwhile, bleach was predicted to be a DNA synthesis inhibitor based on highest similarity to nitrofurantoin, a known inducer of ROS production in bacteria. Interestingly, bleach also correlated to many of the inhibitors of membrane integrity, consistent with bleach imparting generalized oxidative damage on both DNA and membranes across diverse active treatments [33]. Despite the ability to predict MOA by comparing z-scored expression profiles, the PPV of such predictions is lower than from the LOOCV analysis of internal reference data, as evidenced by the overall lower distributions of *r*_*predict*_ values for the external data than the internal data, as well as the lower *r*_*predict*_ values for the same compounds present in both the internal and external datasets (**Fig. 2b**). We attributed these differences to disparities in experimental format, including sublethal dosing that yielded weaker transcriptional signals.

**Fig. 2.**
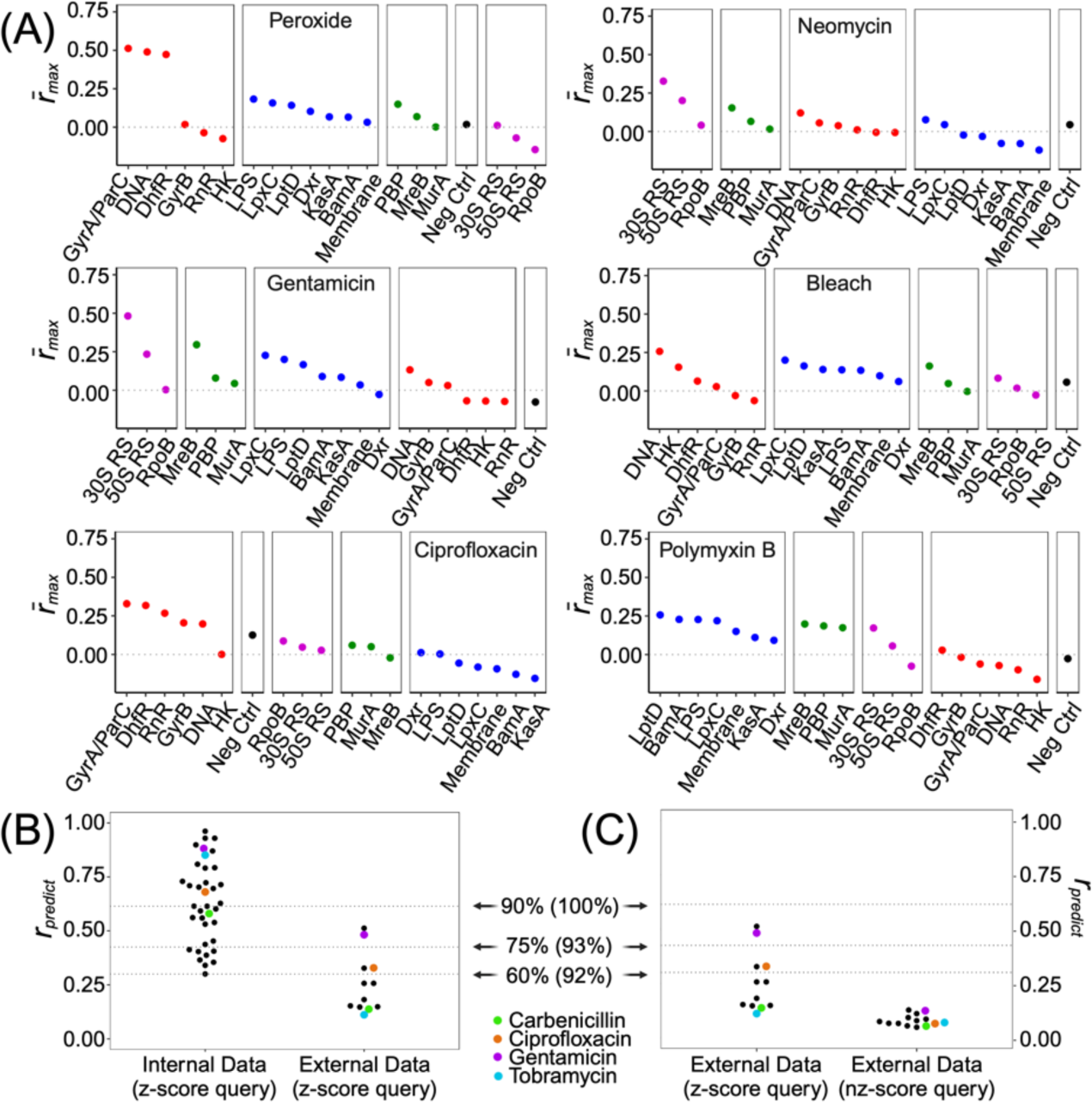
PerSpecTM analysis of external data set by mapping to the antimicrobial reference set. (A) Target correlation scores (*r̅*_*max*_) of six external query compounds against the target categories indicated in the antimicrobial reference set. The highest *r̅*_*max*_ score is the target correlation prediction score (*r*_*predict*_) and identifies the predicted target. Colors correspond to MOA classes: DNA synthesis (red), membrane integrity (blue), cell wall synthesis (green), and protein synthesis (magenta). (B,C) Distribution of *r*_*predict*_ for all compounds in (B) the internal reference set in the LOOCV compared to the external query set using z-scores and (C) the external query set using z-scores compared to nz-scores. Each point represents a compound. Points of the same color identify the same compound in both the reference and external query sets, as shown. Dotted lines indicate three *r*_*predict*_ thresholds, labeled with approximate target PPVs determined from the LOOCV z-score analysis (with MOA PPVs in parentheses). Target prediction scores are not indicated for compounds that were most correlated to negative control.

We next compared the performance of this approach using z-scores calculated across a diverse set of query compound treatments, to using what we termed “nz-scores”, which refers to standardizing gene expression relative only to negative control samples. Specifically, we compared the distribution of *r*_*predict*_ values of the external data using z-scores, calculated across the diverse sample set, versus nz-scores calculated only by comparing to negative controls. Not only was the *r*_*predict*_ distribution of the nz-score analysis much lower, but the nz-score analysis failed to make any meaningful confident predictions (**Fig. 2c**). Indeed, in a similar analysis, nz-scores calculated from within our internal antibiotic reference set, when used as query data, also had a lower distribution of *r*_*predict*_ values and lower confidence in target predictions than when using z-scores, albeit to a lesser extent than for the external data set (**SI Appendix, Fig. S5**). Of note, the external data set only included two negative control samples, which could in part account for the poor performance using nz-scores. Taken together, these results demonstrate that by calculating z-scores across a sample set representing diverse mechanisms, PerSpecTM can yield accurate MOA and target predictions despite highly divergent methodologies. In the absence of sufficient numbers and diversity in a query set, it can be challenging to make any confident predictions, all the more so when divergent experimental methodologies are used which generate weak transcriptional responses.

### MOA and target determination of new antipseudomonal compounds using PerSpecTM and the antimicrobial reference set

To test whether PerSpecTM could predict MOAs of new molecules, we focused on three small molecules we had previously identified using a multiplexed chemical genetic based strategy termed PROSPECT (**PR**imary screening **O**f **S**trains to **P**rioritize **E**xpanded **C**hemistry and **T**argets). PROSPECT is an antimicrobial discovery platform that identifies active small molecules based on their activity against one or a few bacterial hypomorphic mutants, each depleted of a different essential protein target [38]. All three compounds were discovered based on their specific activity against certain hypomorphs in which we had depleted essential outer membrane targets of PA14: PA-0918 against the *lptD-*hypomorph; PA-69180 against *oprL*-hypomorph; and PA-5750 against *lptE*-hypomorph (**Table S5).** LptD and LptE form the LPS transport complex that shuttles LPS across the outer membrane, while OprL is a scaffolding protein involved in cell division [39, 40]. In the case of PA-0918 and PA-5750, we performed expression profiling using PA14 because both compounds were active against wildtype. PA-69180 had no wildtype activity however, and therefore we performed gene expression profiling with the *oprL*-hypomorph.

PerSpecTM predicted PA-0918 to be an inhibitor of membrane integrity with LPS as the predicted target based on an *r*_*predict*_ of 0.74 (100% target PPV, 100% MOA PPV) (**Table S4**; **Fig. 3a**). PA-0918 was most highly correlated with colistin, a polymyxin antibiotic known to target LPS along the outer membrane of Gram-negative bacteria [41]. To test whether PA-0918 shares a similar MOA with colistin as a direct membrane disruptor, we utilized an ethidium bromide uptake assay in which membrane disruption is detected by increased flux of ethidium bromide into the cells. We found that indeed, PA-0918 phenocopies colistin’s ability to promote rapid uptake of ethidium bromide into cells (**Fig. 3b**), consistent with PA-0918 disrupting the outer membrane to increase permeability.

**Fig. 3.**
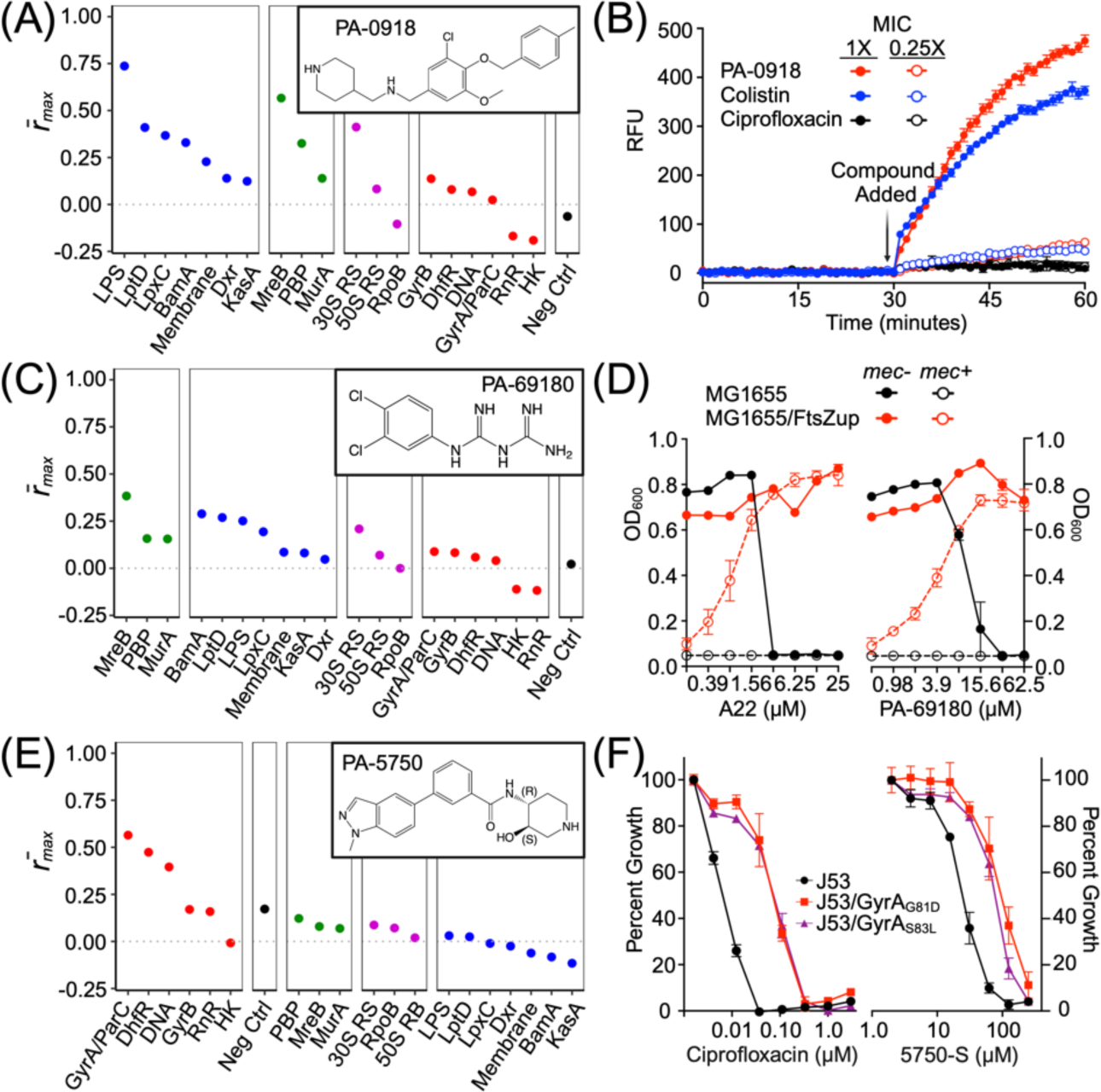
MOA determination of unknown anti-pseudomonal compounds. (A) Target correlation scores (*r̅*_*max*_) of PA14 treated with PA-0918 against the target categories indicated in the antimicrobial reference set. PA-0918 is predicted to be a direct LPS binder, similar to colistin, based on a target correlation prediction score (*r*_*predict*_) of 0.74 (target PPV of 100%). (B) Ethidium bromide uptake in cells treated with PA-0918 and colistin, with ciprofloxacin as a negative control. PA-0918 treatment phenocopies colistin with rapid uptake of ethidium bromide suggestive of membrane disruption. In all experiments, error bars represent S.E.M of three biological replicates (n=3). (C) *r̅*_*max*_ scores of *oprL*-hypomorph treated with PA-69180 against the target categories indicated in the antimicrobial reference set. PA-69180 is predicted to be an MreB inhibitor, similar to A22, based on an *r*_*predict*_ of 0.38 (target PPV of 68%). (D) MG1655 overexpressing *ftsZ* (MG1655/ FtsZup) is resistant to both A22 and PA-69180, and is rescued from mecillinam killing (*mec-* and *mec+* denote 0μg/mL and 2.5μg/mL mecillinam, respectively) by both compounds, implicating MreB as the target. (E) *r̅*_*max*_ scores of PA14 treated with PA-5750 against the target categories indicated in the antimicrobial reference set. PA-5750 is predicted to be a GyrA/ParC inhibitor, similar to ciprofloxacin, based on a *r*_*predict*_ of 0.56 (target PPV 88%). (F) Two ciprofloxacin-resistant *E. coli* J53 mutant strains, each containing single point mutations (G81D or S83L) in GyrA, are cross-resistant to PA-5750, implicating DNA gyrase at the target.

Using the *oprL*-hypomorph, PerSpecTM predicted PA-69180 to be a cell wall synthesis inhibitor with MreB, the major Rod protein responsible for the rod shape of Gram-negative bacteria, as the predicted target based on an *r*_*predict*_ of 0.38 (68% target PPV, 94% MOA PPV) (**Table S4**; **Fig. 3c**). To test whether PA-69180 could be an MreB inhibitor, we used a mecillinam rescue assay described previously [42], which leverages overexpression of the cell-division protein FtsZ in *E. coli* (strain MG1655/FtsZup) to render the Rod system non-essential, thus conferring resistance to MreB inhibition. Meanwhile, MreB inhibition in this strain reverses the lethal effects of the penicillin binding protein 2 inhibitor, mecillinam, which otherwise causes a toxic malfunctioning of the Rod machinery leading to cell death. We found that PA-69180 phenocopies the known MreB inhibitor A22, with MG1655/FtsZup being both resistant to PA-69180 killing and resistant to mecillinam killing upon the addition of PA-69180 (**Fig. 3d**), which was not the case for the negative control ciprofloxacin (**SI Appendix, Fig. S6**). We also generated resistant mutants to PA-69180 in the efflux-deficient strain PAO397 and performed whole-genome sequencing to reveal two mutants, PAO397-r1 and PAO397-r2, revealing mutations at distinct amino acid positions adjacent to the ATP binding pocket of MreB [43] that conferred high-level resistance to PA-69180 and cross-resistance to A22, thereby confirming MreB as the target (**Table S6**).

Finally, PA-5750 was assigned as a DNA synthesis inhibitor with GyrA as the predicted target based on an *r*_*predict*_ value of 0.56 (88% target PPV, 96% MOA PPV) (**Table S4**; **Fig. 3e**). Based on this prediction, we moved to immediately test PA-5750’s ability to both intercalate DNA and inhibit DNA gyrase. We found no evidence of DNA intercalation by PA-5750 based on its inability to stimulate supercoiling activity of topoisomerase I or to displace ethidium bromide from purified plasmid, in contrast to the known DNA intercalator m-amsacrine (m-AMSA) (**SI Appendix**, **Fig. S7a,b**). Instead, we found that PA-5750 inhibited *E. coli* DNA gyrase’s ability to supercoil DNA while the inactive enantiomer of PA-5750 failed to do so (**SI Appendix**, **Fig. S7c**). Lastly, using two ciprofloxacin-resistant mutants of *E. coli* J53 strain, which each harbor different GyrA point mutations, we demonstrated cross-resistance to PA-5750 (**Fig. 3f**), but not to the GyrB inhibitor novobiocin (**SI Appendix, Fig. S8**). Taken together, these results support PerSpecTM’s prediction that PA-5750 targets gyrase, and overall, validate its ability to predict MOA for novel compounds.

### Correlating the transcriptional responses of hypomorphic strains

While we demonstrated the ability of PerSpecTM to predict small molecule MOAs using an antimicrobial reference set, the limited number of known inhibitors and their small set of targets and MOAs could restrict the utility of such an approach, particularly with respect to assigning MOA to compounds with completely novel targets and MOAs. We thus sought to test whether the transcriptional responses of hypomorphic strains, in which essential targets have been depleted, correlate with chemical inhibition of the corresponding targets. If so, PerSpecTM could be used in a reference-based manner to correlate responses between small molecule inhibition to genetic depletion of the same target, thereby providing a comprehensive map of all targetable pathways.

We initially analyzed expression data from the seven hypomorphs we had previously engineered using a regulated promoter replacement strategy to deplete essential gene targets. However, the transcriptional changes in these strains relative to wildtype bacteria were minimal. Based on our findings with small molecules, we wondered whether the transcriptional changes in these strains relative to wildtype bacteria could be minimal due to insufficient strength of knockdown provided by the promoter replacement system. We therefore shifted strategies to use CRISPRi technology to create an inducible gene repression system. We constructed 19 CRISPRi strains encoding arabinose-induced dCas9 and constitutively expressing guide RNAs targeting *dhfR*, *dxr*, *gyrA/gyrA-2* (two different strains with different guides), *gyrB*, *leuS*, *lptA*, *lptD*, *lptE*, *lpxC*, *mreB, murA*, *parC*, *pbpA*, *ssb*, *rplJ*, *rpoB*, *rpsL*, and a no gene control (referred to as CRISPRi-*C26*), summarized in **Table S7** (**SI Appendix, Additional Methods** for details of strain construction). In each CRISPRi strain, the addition of arabinose induced dCas9 expression and subsequent transcriptional repression of the target gene of interest. We characterized the arabinose-dependent growth phenotypes of each CRISPRi strain (**SI Appendix**, **Fig. S9**). The CRISPRi-C26 control strain showed no significant growth delay over an arabinose range of 0–0.5% (w/v), indicating no appreciable dCas9-mediated toxicity in PA14. Mutants targeting *dhfR*, *lptA*, *lpxC*, *mreB*, *murA* and *parC* all demonstrated severe arabinose dose-dependent growth delays, whereas mutants targeting *dxr*, *gyrA*, *gyrB*, *leuS*, *lptD*, *lptE*, *rplJ*, *rpoB* and *rpsL* exhibited more moderate growth delays at higher arabinose concentrations. By comparison, mutants targeting *pbpA* and *ssb* showed no growth delay.

We performed transcriptional profiling of all 19 CRISPRi strains in 384-well format at 360 minutes in 0%, 0.125%, and 0.5% arabinose before chemical lysis, RNA extraction, and RNAseq library construction. Z-scores were calculated across all strains and arabinose doses, thereby accentuating strain-specific transcriptional changes. Additionally, we used DESeq2 to assess the level of knock-down of the targeted genes in each condition, as summarized in **Table S7**. In general, CRISPRi strains demonstrated an arabinose-dependent decrease in the expected target mRNA levels compared to levels in the CRISPRi-*C26* control strain, ranging from 1.5 – 174-fold in 0.125% or 0.5% arabinose for conditions yielding significant on-target gene-expression changes (adjusted p-value < 0.05). Of note, no statistically significant expression changes in the target gene were detected even at the highest arabinose concentrations for mutants carrying guides targeting *lptE*, *ssb* and *rpoB*.

To form a genetic depletion reference set using robust transcriptional profiles from the CRISPRi strains, we compiled all treatment conditions with greater than 0% arabinose. We again removed treatment conditions yielding transcriptionally weak responses, defined as the differential expression between arabinose treatment and control guide C26 wherein no gene had an adjusted p-value that was more significant than the minimum adjusted p-value defined previously by comparing PA14 in water versus DMSO control (i.e. p_adj_ < 10^−20^). Applying this filter, mutants carrying guides targeting *dhfR*, *leuS*, *lptE*, *gyrA-2* (i.e. the CRISPRi-*gyrA-2* strain) were removed from analysis, leaving 14 strains in the genetic depletion reference set, with some comprising multiple levels of knockdown induced by different arabinose doses, in addition to the control strain.

Finally, we applied PerSpecTM using the genetic depletion reference set to assess whether the transcriptional profiles of the CRISPRi strains could serve as a reference. The target correlation scores *r*_*max*_ were calculated between the z-scored gene expression profiles of CRISPRi strains (as the reference set) and those of compound treatments (the query samples) of PA14, except for fosmidomycin, which was tested in the *dxr*-hypomorph strain due to its inactivity against PA14. As shown in **Fig. 4**, the transcriptional responses of antibiotic treatments generally correlated to transcriptional responses of the corresponding CRISPRi strains depleted for the cognate gene targets, or their associated ribosomal/protein complexes. The target was predicted correctly in some cases. Ciprofloxacin treatment correlated strongest with *parC* depletion, followed by *gyrA*, accurately mapping to its known targets of GyrA and ParC. Similarly, the GyrB inhibitor novobiocin correlated most strongly to depletion of *gyrB*, the LpxC inhibitor PF-04753299 mapped to *lpxC* depletion, the LPS transport inhibitor POL7080 mapped to *lptA* depletion, the MurA inhibitor fosfomycin mapped to *murA* depletion, and the aminoglycoside amikacin mapped to ribosomal subunit *rpsL* depletion. Of note, strains depleted for *parC* (a topoisomerase) and *lptA* (part of the LPS transport complex), which showed some of the greatest fitness defects, yielded the strongest *r*_*predict*_ values of 0.56 and 0.46 upon inhibition with ciprofloxacin and POL7080, respectively. When the target was not correctly predicted, in most cases PerSpecTM accurately predicted the MOAs. For example, all protein synthesis inhibitors (amikacin, clarithromycin, and rifampicin) correlated most strongly with the two CRISPRi strains depleted of targets which interfere with protein synthesis (*rpsL*, *rplJ and* r*poB*). Similarly, fosmidomycin, which was misassigned as an LptA inhibitor, was correctly predicted to disrupt membrane integrity. Indeed, the role of its target Dxr in the synthesis of isoprenoid precursors is required for maintenance of the cell wall and thus membrane integrity. Of the two compounds with misassigned MOAs altogether, the cell wall synthesis inhibitors A22 and imipenem, both were predicted to be membrane disruptors while correlating to a lesser extent with CRISPRi strains disrupting cell wall synthesis. In general, like the small molecule reference set, there is significant overlap in transcriptional changes between inhibitors of membrane integrity and cell wall synthesis with compounds tending to correlate with gene depletions across both mechanistic classes, suggesting a shared transcriptional response to disruptors of both cellular processes. Together these data demonstrate that the transcriptional response of chemical inhibition of a given target correlates with that of CRISPRi-induced repression of the same target.

**Fig. 4.**
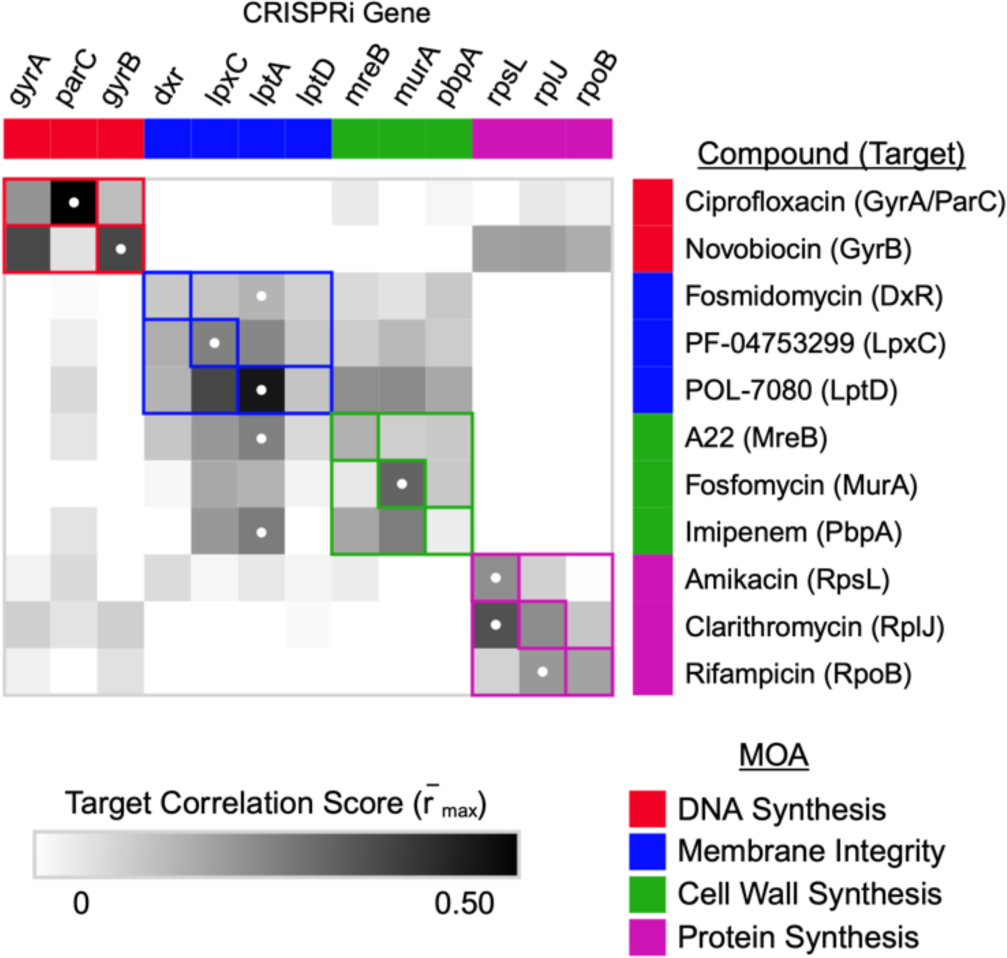
PerSpecTM analysis of antimicrobial treatments queried against the genetic depletion reference set. Heatmap intensities show target correlation scores (*r̅*_*max*_) of 90-minute antibiotic treatments when queried against CRISPRi strains in the genetic depletion reference set treated with 0.125% or 0.5% arabinose to induce dCas9 expression. PA14 was treated with all query compounds except fosmidomycin, which lacked wildtype activity, and thus the *dxr*-hypomorph was used. White dots indicate the target correlation prediction score (*r*_*predict*_) for each query compound. Colored boxes indicate the MOA relating to each target gene or query compound. Note that from the genetic depletion reference set, only those that have corresponding compound treatments targeting the same gene targets, or the associated ribosomal/protein complexes, are shown.

## Discussion

Target identification is central to any antibiotic discovery pipeline to guide candidate prioritization and development. The advancement of more sophisticated approaches to antibiotic discovery is working to expand the chemical space and increase the number of primary candidates discovered with antibacterial activity. Meanwhile, these novel platforms highlight the bottleneck that target identification imposes on antibiotic development efforts. While measuring the transcriptional response to small molecule perturbation has now become quite a standard step in characterizing a new antibiotic candidate, its value in MOA elucidation is often supportive at best, despite the anticipation that it could yield valuable insights into the biology of how target disruption leads to killing, thus leading to a MOA. To date, the interpretation of transcriptional responses has been confounded by background changes in gene expression that are secondary or even unrelated to the immediate, direct effect of target inhibition – such as those relating to growth, general stress, or cell death. These changes can drown out perturbation-specific gene responses. The major approach that has been taken to mitigate these confounders is to tailor experimental designs to control for background gene expression by treating with subinhibitory concentrations while carefully trying to match growth rates; these approaches, however, limit the strength of the elicited transcriptional responses. An alternative approach has been to develop computational algorithms that select the most informative genes, which in some implementations may be at risk of overfitting the reference dataset if the selected genes are optimized based on their ability to predict the MOAs of the compounds in the reference set.

Here we developed a novel strategy, PerSpecTM, that works by leveraging the ability to scale the measurement of expression profiles across a large set of active, diverse perturbations, thereby enabling a standardization (calculation of z-scores) that deemphasizes the background changes to highlight perturbation-specific changes as the basis for more accurate target and MOA prediction. We demonstrate proof-of-principle that by applying PerSpecTM’s reference-based approach, either to a set of known antimicrobial compounds with known MOA or a set of strains which are genetically engineered to be depleted of different essential targets, a compound’s MOA and even target can be reasonably predicted. We validated PerSpecTM’s performance on both internal and external small molecule datasets relative to an antimicrobial reference set, as well as proof-of-principle using a collection of 14 CRISPRi depletion strains.

Importantly, we found that eliciting a strong transcriptional response was critical to PerSpecTM’s ability to predict MOA in the case of both chemical and genetic perturbations. Reasonably potent antimicrobial activity (*i.e.,* being able to treat with above-MIC concentrations) was necessary to ensure a robust transcriptional response to chemical treatments; this however poses a barrier for early MOA prediction of new compounds that have not yet been optimized to have potent wildtype activity. For weakly active molecules, their solubility limits could prevent treatment of wildtype bacteria at 2–4-fold above the MIC. Meanwhile, such weak inhibitors could be of significant value either as early antibiotic candidates or as chemical biological probes, particularly if they work by a novel MOA. In such cases, we show that hypersusceptible strains can be substituted for wildtype bacteria, with the changes in such strains mapping to the same changes in wildtype bacteria, dose adjusted. Thus, engineered, hypersensitive hypomorphs can enable PerSpecTM ‘s ability to predict MOA in the case of weak inhibitors. Meanwhile, we also found that not all genetically engineered hypomorphs have transcriptional profiles that differ significantly from those of wildtype bacteria. Sufficiently strong knockdown, for example in the engineered CRISPRi strains versus the promoter replacement strains, is critical to eliciting a robust transcriptional difference between mutant and wildtype bacteria, even while the level of knockdown required to generate these differing responses will vary from gene to gene. This is supported by the fact that the least fit CRISPRi strains – with growth impairment reflecting sufficient target depletion to impact cellular physiology – yielded the strongest Pearson correlations to chemical inhibition of their corresponding gene targets. Thus, the more impaired a strain’s fitness is, the more informative its transcriptional profile can be.

We applied PerSpecTM to predict MOA for three new active anti-pseudomonal compounds that we had identified – PA-0918, PA-69180 and PA-5750 – to highlight its power in MOA determination early in the antibiotic discovery process. All three compounds originated directly from chemical libraries, and therefore had not yet undergone extensive structure-activity-relationship studies or optimization. PerSpecTM predicted PA-0918 to be a membrane disruptor, PA-69180 to be an MreB inhibitor, and PA-5750 to be an inhibitor of DNA gyrase, all of which we subsequently confirmed with supporting experimental validation. The case of PA-69180, which lacked significant wildtype activity, highlights the advantage of leveraging the hypersusceptibility of the *oprL*-hypomorph to generate a robust transcriptional signal, which enabled a correct, high-confidence MOA prediction. Interestingly, it also suggests a potential functional interaction between OprL and MreB, relating cell shape and structure to cell wall integrity. The case of PA-5750 adeptly highlights the advantage of integrating MOA prediction tools early in the discovery process, as PerSpecTM’s prediction immediately focused our efforts on the hypothesis that PA-5750 inhibits DNA synthesis. Taken together, these data lend credence to the ability of transcriptional profiling to expedite the incorporation of MOA prediction into antibiotic discovery processes, aiding in prioritization before any chemical optimization.

Referenced-based MOA prediction by expression profile mapping to a collection of known antimicrobial compounds is inherently limited by the diversity of the reference set, which poses a barrier to discovering inhibitors of novel biological pathways without known chemical inhibitors to anchor predictions. We therefore explored a complementary approach of mapping to genetic perturbations by constructing a transcriptionally repressed reference set using CRISPRi technology. While chemical inhibition of a series of gene targets correlated strongest with their corresponding CRISPRi knockdown strains and the best correlations were made when the level of knockdown was strong enough to impair fitness, some strains without impaired fitness nevertheless provided informative expression changes. In future iterations of this approach, generating strains in which bacterial fitness is compromised as part of a larger, more comprehensive library of CRISPRi strains, potentially guided by genome-wide fitness studies as have been reported for *M. tuberculosis* [44], would yield a stronger genetic perturbation reference set from which one can generate more robust correlations for MOA prediction. Nevertheless, these data provide proof-of-concept that a dedicated genetic perturbation reference set could facilitate accurate MOA prediction, including for gene targets without known chemical inhibitors. In theory, a comprehensive library of CRISPRi strains could be used to define the transcriptional map of all essential genes of a certain bacterial species, serving as a reference set for completely unbiased MOA predictions.

Nevertheless, several caveats in the PerSpecTM approach warrant comment. PerSpecTM remains a reference-based strategy; therefore, by relying on an antimicrobial reference set as the basis for MOA prediction, PerSpecTM will inherently be limited to predicting MOA for novel compounds which work by known, established mechanisms or targets for which there are pre-existing inhibitors. This limitation can be circumvented by constructing reference sets of hypomorphic mutants, based on the assumption that gene depletion phenocopies target inhibition. While this assumption is often true, it is not universally so. For example, differences in the timescale of loss of function between small molecule inhibition and gene depletion can yield different transcriptional responses as cells adapt with differing compensatory mechanisms. Alternatively, complete absence of a protein may not phenocopy inhibition of a particular protein function, particularly when a protein target has multiple functions.

Relying on a genetic depletion reference set as the basis for MOA prediction also has limitations. Firstly, as not all antibacterial compounds work via a protein target depletion *(e.g.,* membrane disruptors, DNA intercalators, free radical producers), and transcriptional response of depleted essential protein targets may not always phenocopy the response from queried compounds. Secondly, the specific cellular response to some small molecule or genetic perturbations may not be well reflected in strong, canonical transcriptional responses, instead resulting in other changes (*e.g.,* metabolic or structural changes that lead to death). β-lactams are such a case where fewer changes in gene expression occur on a relatively slow timescale compared to other antibiotic classes. Finally, whether using an antimicrobial or genetic depletion reference data set, the MOA prediction resolution of PerSpecTM can be compromised by the complexity of cellular networks which have significant interactions and cross-talk, as evidenced by the overlapping transcriptional profiles of disruption of the cell wall and membrane integrity; inhibition of targets which share common functions or that are highly interactive can generate similar, overlapping transcriptional responses. Meanwhile, off-target effects of small molecules, particularly of weakly potent ones, can similarly generate complex, noisy transcriptional programs that are uninterpretable and do not map to inhibitors of single targets or single gene depletions.

At the same time, we note an important caveat to PerSpecTM’s z-score based approach. The power of the z-score metric is both in its simplicity as well as its ability to emphasize the informative, relevant signal while deemphasizing the noise that is common to all death-inducing perturbations. This work demonstrates the reasonably good performance of such an approach. However, the ability to statistically remove the common noise without losing informative signal in this manner relies on the unbiased representation of a balanced, diverse, distinct set of perturbations analyzed within a batch. Biased MOA representation within the batch might allow a single MOA to dominate, resulting in a z-score analysis dampening signals that are common to, but also potentially meaningful across, a large number of samples. In this work, we tried to minimize bias by ensuring that minimally, each batch of queried samples included control perturbations resulting from inhibition of DNA synthesis, protein synthesis, and either cell wall or membrane integrity (or both), along with negative control samples, which could be used to calculate z-scores. With this representation, PerSpecTM performed well for these datasets. This included an external data set with slightly different MOA composition than the antimicrobial reference set. However, further work would be needed to determine whether the set of control inhibitors we used are sufficiently balanced and unbiased, such that analysis of data using a different albeit still diverse set of controls in a new query data set would replicate this MOA prediction accuracy. Finally, further effort to optimize standardization with respect to negative controls (nz-scores) would be needed to enable accurate prediction as an alternative application of PerSpecTM, without the need for balanced representation of diverse control inhibitors in a given batch.

Here we present a strategy for increasing the ability for transcription-based mapping to predict MOA as well as to increase confidence in predictions with the goal of elevating gene expression profiling to be a driver for primary MOA prediction rather than just as an approach to provide ancillary data supportive of an MOA. As such, in a reference-based manner, the expression response to a small molecule perturbation can serve as a phenotype, and can be used in conjunction with other phenotypic measurements, including microscopy [45–47], chemical-genetic interaction profiles [38, 48, 49], and phenomic profiling using gene deletion libraries or transposon libraries for network mapping of gene interactions in response to chemical treatments for MOA prediction [9, 50, 51]. As we look to the future, the development of a comprehensive suite of complementary phenotypic measurements resulting from small molecule perturbation will hopefully enable more rapid and efficient MOA elucidation, thereby accelerating the discovery, prioritization, and development of novel antibiotic candidates to address the growing problem of antibiotic resistance.

## Materials and Methods

### Bacterial strain construction

*P. aeruginosa* strains were derived from the UCBPP-PA14 lab strain described previously [52]. **Table S8**, **Table S9** and **Table S10** describe all plasmids, strains and primers used in this study. Detailed methods of strain construction are available in the **Additional Methods** of **SI Appendix**.

### Gene expression experiments and library construction

Gene expression experiments were carried out in biological triplicate in 384-well microplates (Thermo Fisher Scientific, Waltham MA). Stock solutions of compounds were dissolved fresh in DMSO or water, and 30μL of compound solutions were mixed with 30μL of mid-log culture to yield 1×10^8^ CFU/mL bacteria and compounds in 0.5% (v/v) water (or DMSO) in LB. Cells were treated at 37°C without shaking in a humidity chamber for 30, 60, 90, and 120 minutes. In the case of the CRISPRi strains, cells were treated for 360 minutes. At the end of the incubation period, samples were then mixed with 30μL of 3X RNAgem Blue Buffer (Custom Science, New Zealand) and chemically lysed by incubation in a thermocycler at 75°C for 10 minutes. Total RNA was extracted using the Direct-zol kit (Zymo Research, Irvin CA), and RNA quality and quantity were analyzed using the RNA ScreenTape with the 2200 TapeStation (Agilent, Santa Clara CA). Illumina cDNA libraries were generated using a modified version of the RNAtag-seq protocol [53], as outlined in more detail in the **Additional Methods** of the **SI Appendix**.

### Computational analysis of gene expression profiling

#### Processing raw reads

Sequencing reads from each sample in a pool were demultiplexed based on their associated barcode sequence using custom scripts, allowing up to 1 mismatch in the barcode. Barcode sequences were removed from the first read as were terminal G’s from the second read that may have been added by SMARTScribe during template switching. Sequencing reads were aligned to the UCBPP-PA14 reference sequence using BWA [54], assigned genomic features using custom scripts, and visualized using GenomeView [55]. Samples that had low read counts, where over 80% of the genes had zero counts, were filtered out of the dataset. In addition, any genes that had zero counts in 50% or more of the samples in a sequencing batch were removed.

#### Assessing robustness of transcriptional response

DESeq2 [56] was used to calculate the log_2_ fold change and adjusted p-values (with target false discovery rate of 0.05) in gene expression to treatment relative to vehicle control, as well as to DMSO versus water control. We used the comparison of gene expression between DMSO and water as a baseline for setting a threshold to separate “weak” from “robust” responses. For a transcriptional response to be considered robust, the gene expression of at least one gene had to be different from the negative control with an adjusted p-value that is smaller than 10^−20^. Conditions with weak transcriptional responses were excluded from the reference set, as were conditions with transcriptional responses more highly correlated to negative control than any other condition in the dataset, including other time points or doses of the same compound.

#### Calculation of moderated z-scores and nz-scores

Read counts were transformed using the variance stabilizing transformation from the DESeq2 package (v1.28.1) in R (4.0.0) [56]. Then, to identify gene expression changes of interest, gene expression values were standardized by calculating moderated z-scores, as described previously [57]. First, the vst-transformed expression values were quantile-normalized across samples in each batch, where each batch was defined as all samples on the same 384-well plate sharing the same strain and time point. Then, robust z-scores for each gene were calculated per batch as follows:

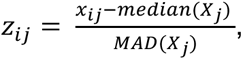

where *i* is the sample, *j* is the gene, *x_ij_* is the quantile-normalized gene expression value for gene *j* in sample *i*, *X_j_* is the vector of all quantile-normalized gene expression values for gene *j* in all the samples in the batch, and MAD is the median absolute deviation calculated using the stats package in R with the default constant of 1.4826 [57]. In cases where there was not a diverse set of mechanisms represented by the compounds in the batch, expression levels were standardized using the median and MADs of only the negative control samples from each batch, rather than all the treatments in the batch, termed “nz-scores” (z-scores relative to negative controls). The datasets that used nz-scores in this manuscript comprised the external dataset, when used to compare with the z-score approach, and the screening hit PA-69180 that was used to treat the *oprL*-hypomorph. All other data involved a diverse set of compounds within each batch, such that gene expression could be standardized across the all the samples in the given batch. Next, replicate data were combined using a weighted average, where the weights were determined by the Spearman correlation between replicate transcriptional profiles, as described previously [57]. Finally, extreme z-score values were clipped such that fewer than 1% of the values were affected. For the reference set, z-score values lower than −10 were set to −10, and those higher than 10 were set to 10, so that extreme values would not spuriously impact downstream correlation values. After calculation of all z-score values, transcriptional data were visualized using a UMAP [58], implemented by the R package UMAP (v0.2.10.0).

#### Selection of marker genes to validate transcriptional responses with known biology

Within each antibiotic class, the z-scored gene responses to 90-minute compound treatment were sorted for strength (i.e. absolute value of z-score) and consistency (i.e. same sign of z-score across all samples). The minimum absolute value z-score was recorded for each gene, across all the compound treatments in each antibiotic class, to capture consistently strong effects within each class. If the z-scores for a given gene had disparate signs within the antibiotic class, that gene was removed from analysis for that class. Top up- and down-regulated genes were assessed for validation with known biology of the targets.

#### Prediction of compound targets and leave-one-out cross validation

The reference set of transcriptional profiles was used to predict the targets of unknown compounds by using Pearson correlation values. Depending on the composition of each query dataset, z-score or nz-scores were calculated. Individual query compound treatments (e.g. multiple doses or time points) were correlated to the reference set z-score values. For each query treatment, the maximal Pearson correlation to all the treatments in the given target category in the reference set was recorded (*r*_*max*_). Any query treatment that was most highly correlated to vehicle control was removed. Then, if there were multiple remaining treatments for the given query compound, the average of their *r*_*max*_ values was calculated to be the target correlation score, *r̅*_*max*_, for each target category. The highest *r̅*_*max*_ value was termed the target correlation prediction score, *r*_*predict*_. The target category corresponding to the *r*_*predict*_ value was then taken to be the predicted target of the query compound, similar to methods described previously [6].

The LOOCV was performed wherein each compound in the reference set was taken to be a query compound. When querying a given compound, all other instances of the same compound (e.g. different doses, time points, or strains) were removed from the reference set. Target predictions were made for each compound, and corresponding MOA predictions were assigned. The PPVs were recorded during the leave-one-out cross validation to be used as a measure of confidence for predictions on unknown compounds. Note that target categories comprised of only one compound (27% of compounds), which could never be predicted correctly in the leave-one-out cross validation, were maintained in the calculation of PPV to obtain a conservative estimate. PPV was calculated both at the level of specific target prediction, and at the level of general MOA prediction. Also, the LOOCV was performed for querying both z-scores and nz-scores, separately, so that PPVs could be assigned for queries using z-scores and nz-scores appropriately. PPVs were calculated using the pROC library (v1.16.2) in R. Predictions with *r*_*predict*_ below the minimum *r*_*predict*_ in the LOOCVs were assigned a PPV of NA, since no confidence could be assessed due to being out of the range of our reference data.

#### Using transcriptional profiles of CRISPRi strains to predict compound mechanisms

Transcriptional data from the CRISPRi strains were processed in the same manner as those from compound treatments using z-score calculations across the batch. Differential gene expression between each sample and the control strain (CRISPRi-C26) at matched doses of arabinose was used to assess the strength of transcriptional response, using the same 10^−20^ p-value threshold to filter out weak signals. Target and MOA predictions of were performed as described previously, but using the CRISPRi transcriptional profiles as the reference data set. Pearson correlation values were calculated between the z-scores of the CRISPRi strains and those of the wildtype strain treated with different compounds targeting the same genes as the CRISPRi constructs. The exception was fosmidomycin, for which the *dxr*-hypomorph strain was used instead of wildtype since the compound was not active against wildtype PA14. A target correlation score, *r̅*_*max*_, was calculated for each pair of compounds (as the query) and CRISPRi strains (as the reference set), as described above. The transcriptional data used as the query were the relevant 90-minute treatments from the antimicrobial reference data set.

#### Scripts and data accessibility

Scripts for calculating z-scores, nz-scores and predicting compound targets were written in R and stored in github (https://github.com/broadinstitute/psa_rnaseq_manuscript_rproject). All raw sequencing data and z-score values can be found on the Gene Expression Omibus (GEO accession GSE251671, https://www.ncbi.nlm.nih.gov/geo/)

### Biochemical characterization of screening hits

Growth kinetics, MIC experiments, resistance selection, as well as assays quantifying ethidium bromide uptake and displacement, mecillinam rescue, DNA intercalation, and DNA gyrase activity, are described in detail in the **Additional Methods** of the **SI Appendix**.

## Acknowledgments

This work was supported by NIH-U19AI142780 (DTH) and NIH-K08AI148581 (KPR), and by a generous gift from Anita and Josh Bekenstein. Gene expression libraries were constructed and sequenced at the Broad Institute of MIT and Harvard by the Microbial ‘Omics Core and Genomics Platform, respectively. We thank Nirmalya Bandyopadhyay, James Gomez, and Roby Bhattacharyya for their helpful discussions. We declare that we have no conflicts of interest. All figures were created with BioRender.com.

