## Supplementary material for "Perturbation-Specific Transcriptional Mapping for unbiased target elucidation of antibiotics": SI Appendix

**This PDF file includes:**

Additional Methods
Figures S1 to S9
Tables S1 to S10
SI References

### Additional Methods

#### Hypomorphic strain construction

**Table S8**, **Table S9** and **Table S10** contain information about all plasmids, strains and primers used in this study. Five hypomorphs (*murA*, *gyrA*, *lpxC*, *dhfR*, or *bamA)*, which encoded alleles under the control of a fixed, weakened promoter, were reported previously [1]. The *dxr, lptE, oprL* and *lptD* hypomorphs were constructed to encode an allele under the control of the $P_{araBAD}$ promoter system described previously [2]. To create the arabinose-controlled hypomorphic strains (except the *lptD*-hypomorph), pEXG2-AmpR knockout plasmids [3] were constructed via 4-part Gibson Assembly (New England Biolabs, Ipswich MA) to position the kanamycin cassette flanked by the upstream and downstream regions of the genes-of-interest. The four pieces included: (1) the upstream region (specific primer information in **Table S10**); (2) the downstream region (specific primer information **Table S10**); (3) the kanamycin resistance cassette (primer pair 21 and 22); and (4) the pEXG2-AmpR backbone (primer pair 19 and 20). Merodiploids were then generated via homologous recombination on cellulose membranes (Sartorius Stedim Biotech, Goettingen Germany) by spotting 20μL of a 1:2:2 mixture of the recipient PA14 strain, 10-beta cells harboring the desired pEXG2-AmpR knockout plasmid, and helper strain pRK2013 [4]. After mating at 37°C for 10 hours, cells were resuspended in 400μL LB and merodiploids were selected on LB agar with 10 µg/mL irgasan (Goldbio) and 200 µg/mL carbenicillin (Invitrogen) and confirmed by sequencing. Gene copies were then introduced into each merodiploid at the neutral *attB*-Tn7 site as previously described [5]. To this end, promoters and genes-of-interest were first cloned via 3-part Gibson Assembly into the pUC18T-miniTn7T-Gm suicide plasmid, obtained as a gift from Herbert Schweizer [5]. The three pieces included; (1) the gene-of-interest (specific primer information in **Table S10**); (2) the $P_{araBAD}$ promoter (primer pair 9 and 10); and (3) the pUC18T-miniTn7T backbone (primer pair 23 and 24). All bacterial conjugations were performed on cellulose membranes by spotting 20μL of a 1:2:2:2 mixture of the recipient merodiploid strain for given gene-of-interest, 10-beta cells harboring the corresponding pUC18T-miniTn7T-Gm plasmid, and helper strains pRK2013 and pTNS3. After mating at 37°C for 10 hours, cells were resuspended in 400μL LB and selected on LB agar containing 30μg/mL gentamicin and 10μg/mL irgasan, permitting growth of only merodiploids encoding the integrated pUC18T-miniTn7T-Gm plasmid. Finally, the second step of homologous recombination was performed for all strains to substitute the native copy of each gene-of-interest with the kanamycin-resistance cassette carried on the pEXG2-AmpR plasmid. Bacteria were incubated in 400μL LB for 2 hours, and subsequently streaked on LB agar plates with 10% sucrose and 200 µg/mL kanamycin. Final colonies were isolated and confirmed by sequencing.

In the case of the *lptD*-hypomorph, the $P_{araBAD}$ promoter, upstream region (primer pair 5 and 6), and downstream region (primer pair 7 and 8) of the native *lptD* operon were first amplified individually, stitched together into one single product, and subsequently integrated into the pEXG2-AmpR knockout plasmid via gateway cloning (Invitrogen, Carlsbad CA). Two step-homologous recombination was subsequently performed as described above, effectively replacing the native promoter of the *lptD* operon with the $P_{araBAD}$ promoter without additional antibiotic selection markers. All clones were confirmed by sequencing.

**CRISPRi strain construction**

All sgRNA sequences and primer information are shown in **Table S7** and **Table S10**. Briefly, we first modified the pUC18T-miniTn7T-Gm (pUC18-derived mini-Tn7 integration vector with gentamicin cassette and $P_{araBAD}$ promoter system) encoding *pmrB*, described previously by our group [6]. First, three *BsmBI* restriction sites were removed site by QuikChange XL mutagenesis kit (Agilent). Next, the Spas-handle gene block (**Table S10**) was purchased (Integrated DNA Technologies) containing the following features: (1) 5' HindIII restriction site; (2) *E. coli* ProD promoter; (3) two unique BsmBI restriction sites immediately 5' to the sgRNA scaffold sequence; (4) rrnB T1 terminator; and (5) KpnI restriction site. The Spas-handle gene block and modified pUC18T-miniTn7T-Gm plasmid were then double-digested with *HindIII* and *KpnI* and ligated with *T4 DNA Ligase* (New England Biolabs, Ipswich MA), effectively replacing the terminator region downstream of the *pmrB* gene with the sgRNA promoter, *BsmBI* cut sites, scaffold, and terminator. The dead CRISPR-associated protein 9 (dCas9) from *Streptococcus pasteurianus* (*Spas*) was purchased from Addgene (Watertown, MA), and assembled with the pUC18T-miniTn7T-Gm with sgRNA handle via 2-piece Gibson Assembly (primers 33 and 24; primers 31 and 32), inserting a c-terminal his6 tag in the process. Finally, complementary sgRNA targeting oligos (**Table S5**) were then annealed and ligated (T4 DNA ligase, (New England Biolabs) into the BsmBI-digested CRISPRi vector backbone using NEBridge Golden Gate Assembly (New England Biolabs, Ipswich MA).

**Gene expression library construction**

Illumina cDNA libraries were generated using a modified version of the RNAtag-seq protocol [7]. 250ng of extracted RNA was fragmented, dephosphorylated, and ligated to DNA adapters carrying 5'-$AN_8$-3' barcodes of known sequence with a 5' phosphate and a 3' blocking group. Barcoded RNAs were pooled and depleted of rRNA using the Pan-Bacteria riboPOOL depletion kit (siTOOLs Biotech, Galen Laboratories). Pools of barcoded RNAs were converted to Illumina cDNA libraries in 2 main steps: (i) reverse transcription of the RNA using a primer designed to the constant region of the barcoded adaptor with addition of an adapter to the 3' end of the cDNA by template switching using SMARTScribe (Clontech) as described [8]; (ii) PCR amplification using primers whose 5' ends target the constant regions of the 3' or 5' adaptors and whose 3' ends contain the full Illumina P5 or P7 sequences. cDNA libraries were sequenced on the Illumina [NovaSeq SP 100] platform to generate paired end reads.

**MIC determination and growth kinetics**

All MIC and growth kinetic experiments were conducted in biological triplicate in 384-well microplates (Thermo Fisher Scientific, Waltham MA) using the broth microdilution method previously described [9]. All arabinose-inducible hypomorphic strains were grown initially in the presence of 0.25% (v/v) arabinose in LB, and subsequently sub-cultured without arabinose. Growth curves and MIC experiments of hypomorphic strains were performed without arabinose by back-diluting mid-log cultures to $5x10^5$ CFU/mL bacteria ($OD_{600}$ 1.0 = $10^9$ CFU/mL based on plate counts). Growth curves for all CRISPRi strains were performed at various doses of arabinose, the inducer of dCas9. For MIC assays at high inoculum, mid-log bacterial cultures were diluted to 1 x $10^8$ CFU/mL bacteria ($OD_{600}$ 1.0 = $10^9$ CFU/mL based on plate counts). Antimicrobial compounds were serially diluted in either DMSO or water to a final concentration of 0.5% (v/v) in LB. For MIC experiments, microplates were incubated in a humidity chamber without shaking for 16 hours at 37°C, after which time the $OD_{600}$ was measured in the Spark Multimode Reader (Tecan, Männedorf Switzerland). For growth kinetics, Microplates were incubated in a large humidity cassette in the Spark Multimode Reader (Tecan, Männedorf Switzerland) for 24 hours at 37°C, and $OD_{600}$ measurements were recorded after 30-seconds of shaking every hour. Data were plotted and analyzed using GraphPad Prism10 (GraphPad Software, Boston MA) to determine growth kinetic curves or MICs, the latter defined as the minimal compound concentrations required for complete inhibition of bacterial growth.

**Ethidium bromide uptake assay**

PA14 was grown in LB to mid-log phase, washed in PBS, and resuspended to a final optical density of 0.8 in PBS. 50µL of resuspended cells were added to a black, clear bottom 96-well plate (Thermo Fisher Scientific, Waltham MA) with 10µL of 10µM ethidium bromide and 30µL of PBS, making the final bacterial inoculum approximately $4x10^8$ CFU per well. Fluorescence (Ex/Em 530/585) was measured every 30 seconds using an Infinite F200 plate reader (Tecan, Männedorf Switzerland). After 30 minutes of incubation at 37°C, 10µL of compound prepared at 10X final concentration or PBS were added to each well. Colistin and ciprofloxacin were used as positive and negative controls, respectively, dosed at 4-fold and 1-fold their MICs. Fluorescence was measured every 30 seconds for 90 minutes at 37 °C. Non-intercalated ethidium bromide fluorescence was removed using bacteria-free wells. The change in fluorescence from compound treatment was plotted using GraphPad Prism10 software relative to untreated PA14.

**Mecillinam rescue assay**

Mecillinam rescue experiments were carried out in biological triplicate as described previously [10]. Briefly, in 384-well microplates (Thermo Fisher Scientific, Waltham MA), 20µL of mid-log *E. coli* MG1655 culture (or MG1655/FtsZup) were mixed with 30µL compound solution, yielding a final bacterial inoculum of 200 CFU in LB with 0.5% (v/v) DMSO. After plate incubation at room temperature for 30 minutes, 10µL of 15µg/mL mecillinam solution was added to create a final condition comprising 200 CFU of MG1655 (or MG1655/FtsZup), 2.5µg/mL mecillinam (or LB only), and serially diluted compound (PA-69180, A22 or ciprofloxacin). The microplate was incubated at 30°C without shaking in a humidity chamber for 96 hours. Endpoint $OD_{600}$ measurements were recorded in the Spark Multimode Reader (Tecan, Männedorf Switzerland), and plotted using GraphPad Prism10 software to determine the MICs in the absence of mecillinam, as well as the minimal compound concentration necessary for rescued growth in the presence of mecillinam (Min MIC supp).

**PA-69180 resistance selection on solid agar**

Efflux-deficient PA0397 cells, obtained as a gift from Herbert Schweizer, were grown to mid-log phase in LB and ~$10^8$ CFU were plated onto agar plates containing 50µM PA-69180 (~4X the MIC on LB agar). Resistant mutants were isolated after incubation at 37°C for 16 hours. After re-culturing the colonies in LB with 50µM PA-69180, total DNA was extracted using the DNeasy Blood and Tissue Kit (Qiagen, Hilden Germany). cDNA libraries were constructed as described previously [6]. DNA quality and quantity were determined using the D5000 ScreenTape with the 2200 TapeStation (Agilent, Santa Clara CA). Samples were diluted to a final concentration of 6.8ng/μL and pooled for sequencing on the MiSeq instrument (Illumina, San Diego CA). Annotated reference genomes were obtained from www.pseudomonas.com [11]. Illumina reads were mapped to the PA14 genome and single nucleotide polymorphisms were identified using the Pilon program previously reported [12].

**DNA Intercalation Assay**

DNA intercalation assays were performed with enzymes and reagents from a commercial DNA unwinding kit (Topogen, Buena Vista CO). Briefly, 10X topoisomerase I buffer, pHOT1 supercoiled DNA, and 2 units of topoisomerase I were incubated at 37°C for 30 minutes, generating relaxed DNA substrate. After adding control and test compounds, samples were incubated at 37°C for 30 additional minutes. Reactions were terminated by adding 1% SDS, followed by 50μg/mL Proteinase K with incubation at 56°C for 15 minutes. After preparing the samples in loading buffer, the aqueous phase was extracted using chloroform:isoamyl alcohol (24:1) and loaded onto a 1% (w/v) agarose gel cast in TPE buffer (36mM Tris-HCl, pH 7.8, 1mM EDTA, 30mM $NaH_2PO_4$) and 0.2 μg/mL chloroquine. The gel was electrophorized at 15V for 15 hours, rinsed in distilled water for 15 minutes, stained with 0.5μg/mL ethidium bromide for 30 minutes, and destained in distilled water for 30 minutes. Gel was imaged with a UV transilluminator.

**Ethidium bromide displacement assays**

Serially-diluted ethidium bromide was mixed with an in-house purified plasmid (pHERD-$P_{rhaBAD}$) and compound (m-AMSA, PA-5750 or ciprofloxacin), yielding final conditions with 25–0.2μM ethidium bromide, 2.5ng/μL plasmid DNA, and 100μM, 50μM or 10μM compound. Samples were prepared in a black, clear bottom 96-well plate and equilibrated at 37°C. Fluorescence (Ex/Em 530/585) was measured at until a stable endpoint was reached. Endpoint fluorescence was plotted against ethidium bromide concentration and fit with a one site binding regression using GraphPad Prism10 software.

**DNA gyrase assay**

DNA gyrase assays were performed with enzymes and reagents from a commercial *E.coli* DNA gyrase screening kit (Topogen, Buena Vista CO). Briefly, 5X assay buffer, 250ng of pHOT1 relaxed DNA, 2 units of gyrase, and test compounds (or vehicle control) were incubated at 37°C for 60 minutes. Reactions were terminated by adding 1% SDS, followed by 50μg/mL Proteinase K with incubation at 56°C for 15 minutes. Samples were prepared in loading buffer and the aqueous phase extracted, as described for the DNA intercalation assay. Samples were loaded onto a 1% (w/v) agarose gel cast in TAE buffer (40mM Tris-base, pH 8.3, 20mM acetic acid, 1 mM EDTA, 30 mM $NaH_2PO_4$). The gel was electrophorized at 50V until dye has traveled 60% down length, rinsed in distilled water for 15 minutes, stained with 0.5μg/mL ethidium bromide for 30 minutes, and destained in distilled water for 10 minutes. Gel was imaged with a UV transilluminator.

**Supplemental Figures**

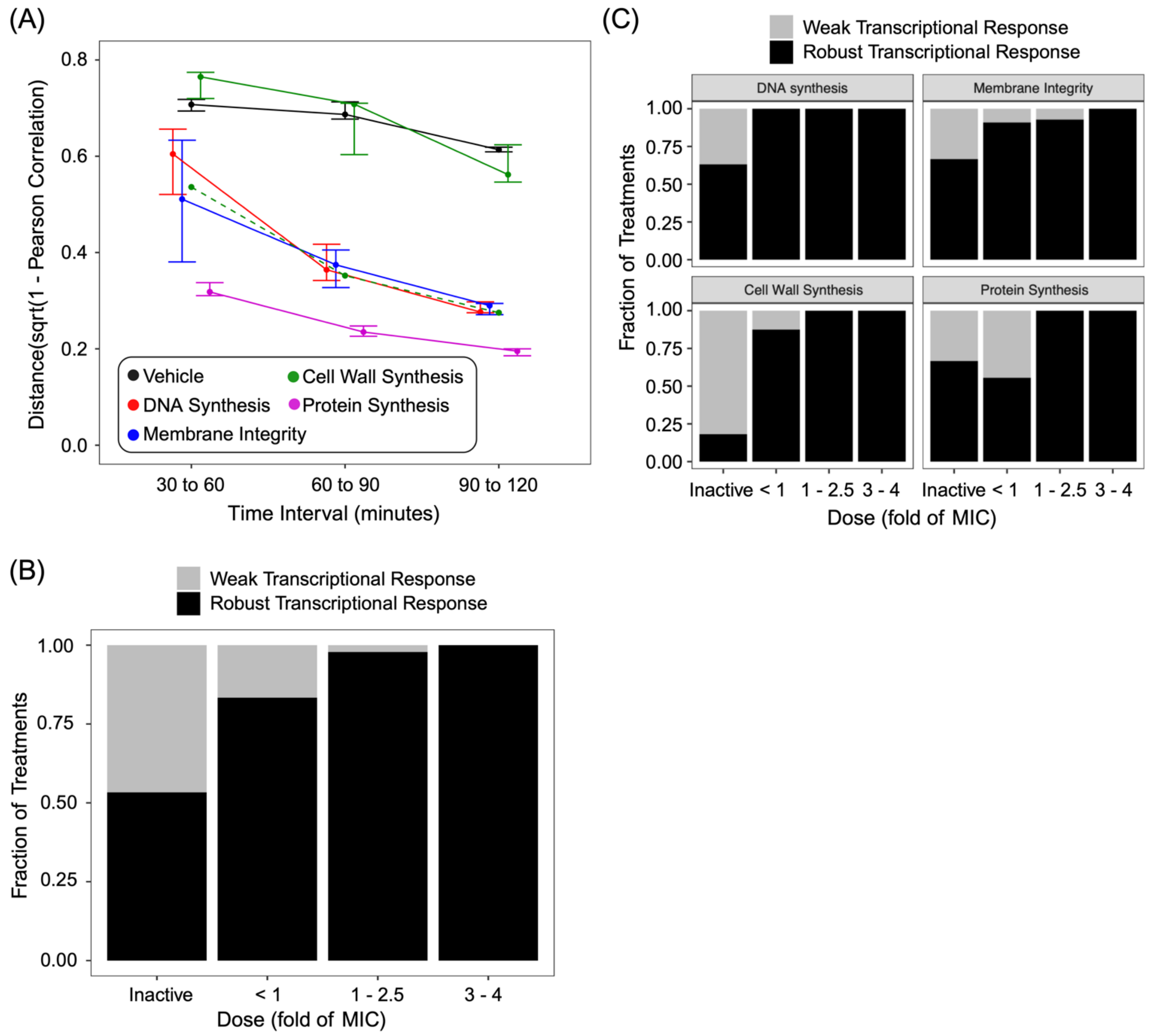


**Fig. S1.** Gene expression time trial and dose response. (A) Pearson distance ($\sqrt{(1 - Pearson\ Correlation)}$) between adjacent time points plateau to a stable endpoint for inhibitors of DNA synthesis, membrane integrity, and protein synthesis. Vehicle and inhibitors of cell wall synthesis demonstrate more variability throughout the time course. Imipenem (dotted green line), a cell wall synthesis inhibitor which does not exhibit a significant inoculum effect, also plateaued to a stable transcriptional endpoint by 90 minutes. (B) The fraction of all 90-minute treatments having a robust transcriptional response as a function of dose relative to the MIC. (C) The fraction of all 90-minute treatments having a robust transcriptional response as a function of dose relative to the MIC, partitioned by MOA: inhibitors of

DNA synthesis, membrane integrity, cell wall synthesis, and protein synthesis. Note that a robust transcriptional response is defined by the differential expression between compound treatment and vehicle control, where at least one gene has an adjusted p-value that is more significant than the minimum adjusted p-value found when comparing PA14 in water versus DMSO control (i.e. $p_{adj} < 10^{-20}$).

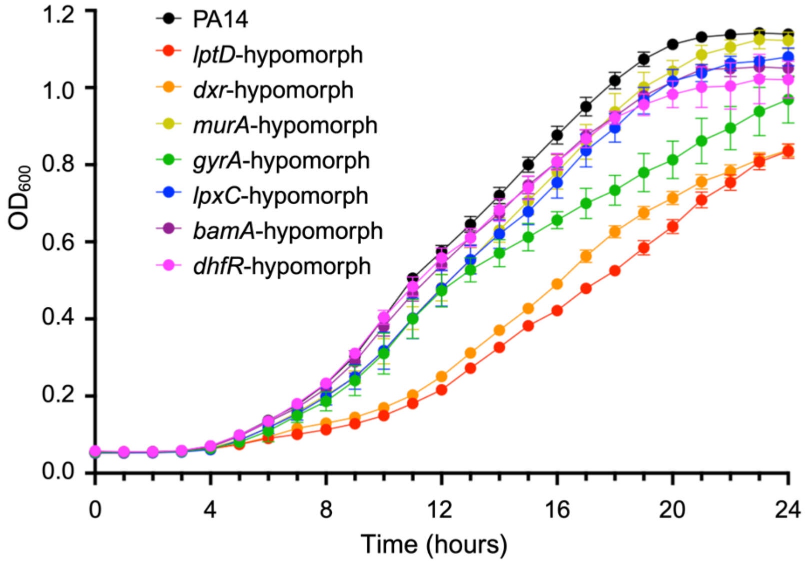


**Fig. S2.** Growth phenotypes of hypomorphs used in this study. The *lptD* and *dxr* hypomorphs exhibited a moderate growth delay compared to PA14. The *gyrA*-hypomorph showed a slight growth delay, and all remaining hypomorphs showed wildtype-like growth.

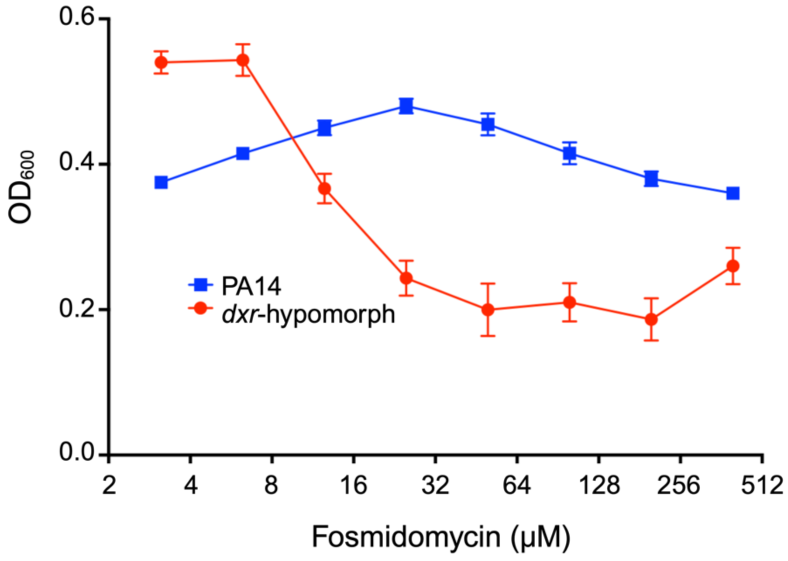


**Fig. S3.** MIC curves of fosmidomycin against PA14 and *dxr*-hypomorph at high inoculum. At high inoculum (starting inoculum $OD_{600}$ 0.1 or 1 x $10^8$ CFU/mL), fosmidomycin had little effect of PA14 growth (blue squares) but induced a significant growth delay of *dxr*-hypomorph (red circles). Error bars represent S.E.M of three biological replicates (n=3).

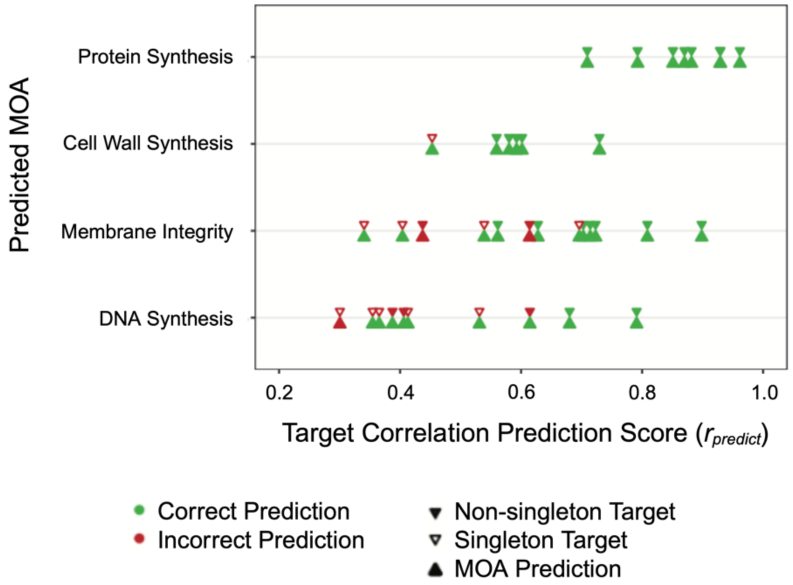


**Fig. S4.** Target correlation prediction scores ($r_{predict}$) from the leave-one-out cross validation (LOOCV) of the antimicrobial reference set, from which positive predictive values (PPVs) are obtained. The relationship between correct predictions of target and MOA relative to $r_{predict}$ in each MOA category is shown. Each pair of triangles correspond to a single compound in the reference set. The triangle above the gray line indicates whether the target for that compound was predicted correctly (green) or incorrectly (red), while the triangle below the gray line indicates whether the MOA for the compound was predicted correctly. Open triangles on top indicate that the given compound is the only representative compound in its target category, and thus had no chance of its target being predicted correctly in the leave-one-out cross validation. PPVs for target and MOA are obtained as the percent of compounds whose target and MOA, respectively, were predicted correctly above a given $r_{predict}$ threshold in the LOOCV. Higher $r_{predict}$ values correspond to a higher percentage of compound targets being predicted correctly, and thus, higher confidence in the prediction. Note that the distribution of $r_{predict}$ values and PPVs vary between different MOA categories.

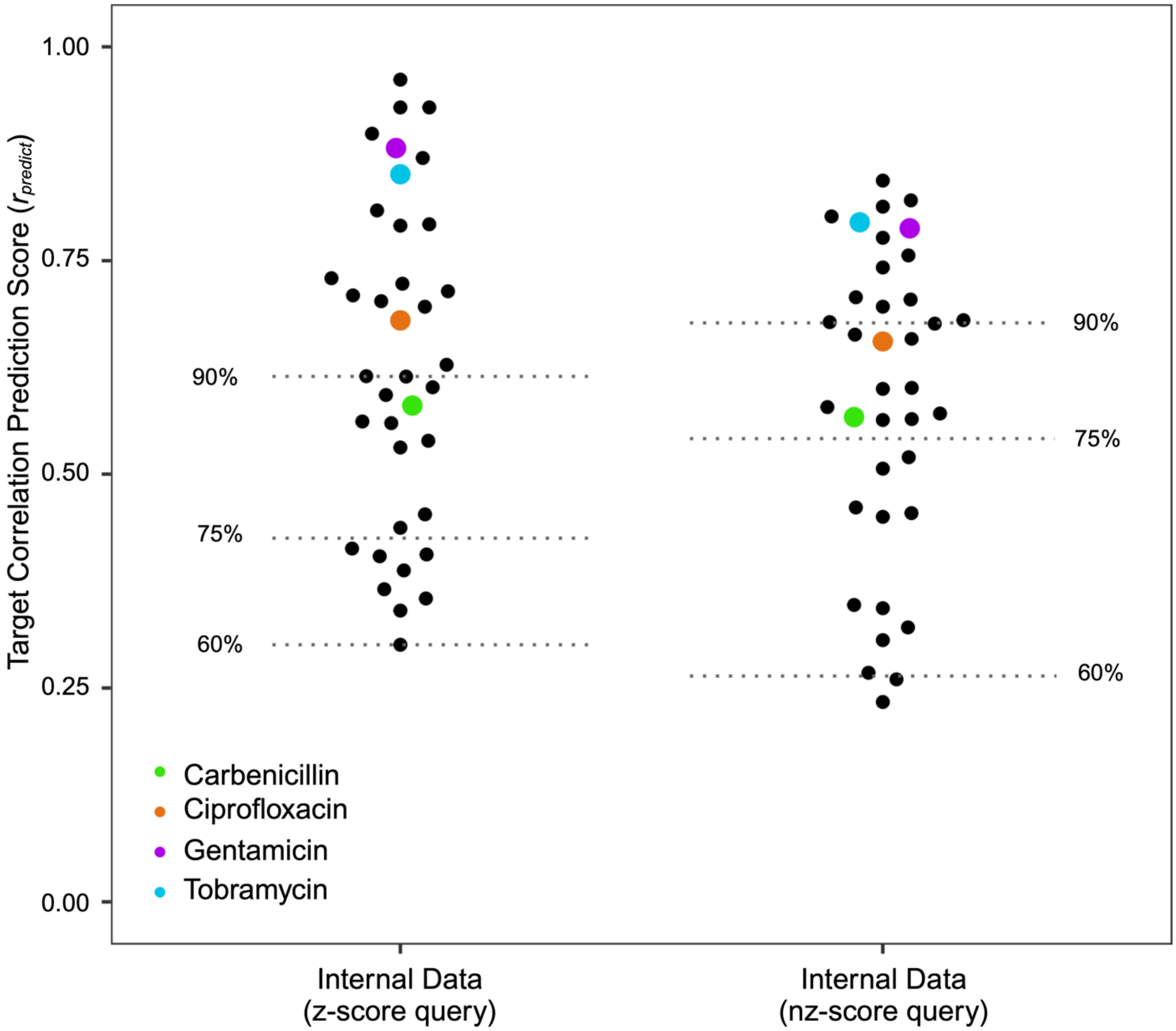


**Fig. S5.** Distributions of $r_{predict}$ for all compounds in the internal reference set in the LOOCV comparing the use of z-scores calculations to nz-score calculations for the query data. Each point represents a compound, and dotted lines indicate three $r_{predict}$ thresholds that were matched by target PPVs in each approach, having approximate PPVs for target prediction labeled. Note that the $r_{predict}$ values arising from nz-score queries are generally lower than those from z-score queries. In addition, the target predictions for the nz-score analysis are generally of lower confidence than for the z-score analysis. For example, more compound predictions fall above the 90% target PPV threshold in the z-score analysis than in the nz-score analysis.

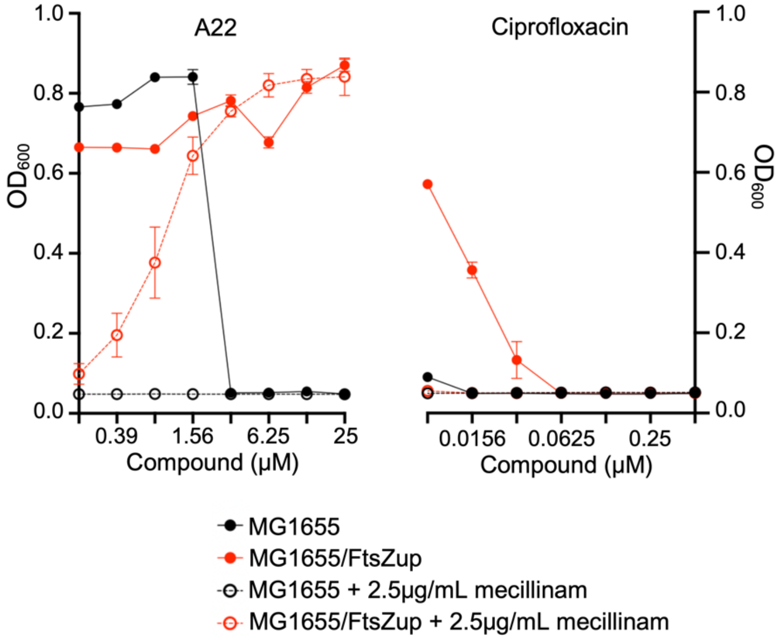


**Fig. S6.** Mecillinam rescue experiments with A22 and ciprofloxacin. MG1655 overexpressing *ftsZ* (MG1655/ FtsZup) exhibits A22 resistance, as well as protection from mecillinam killing by A22, but not for the negative control ciprofloxacin.

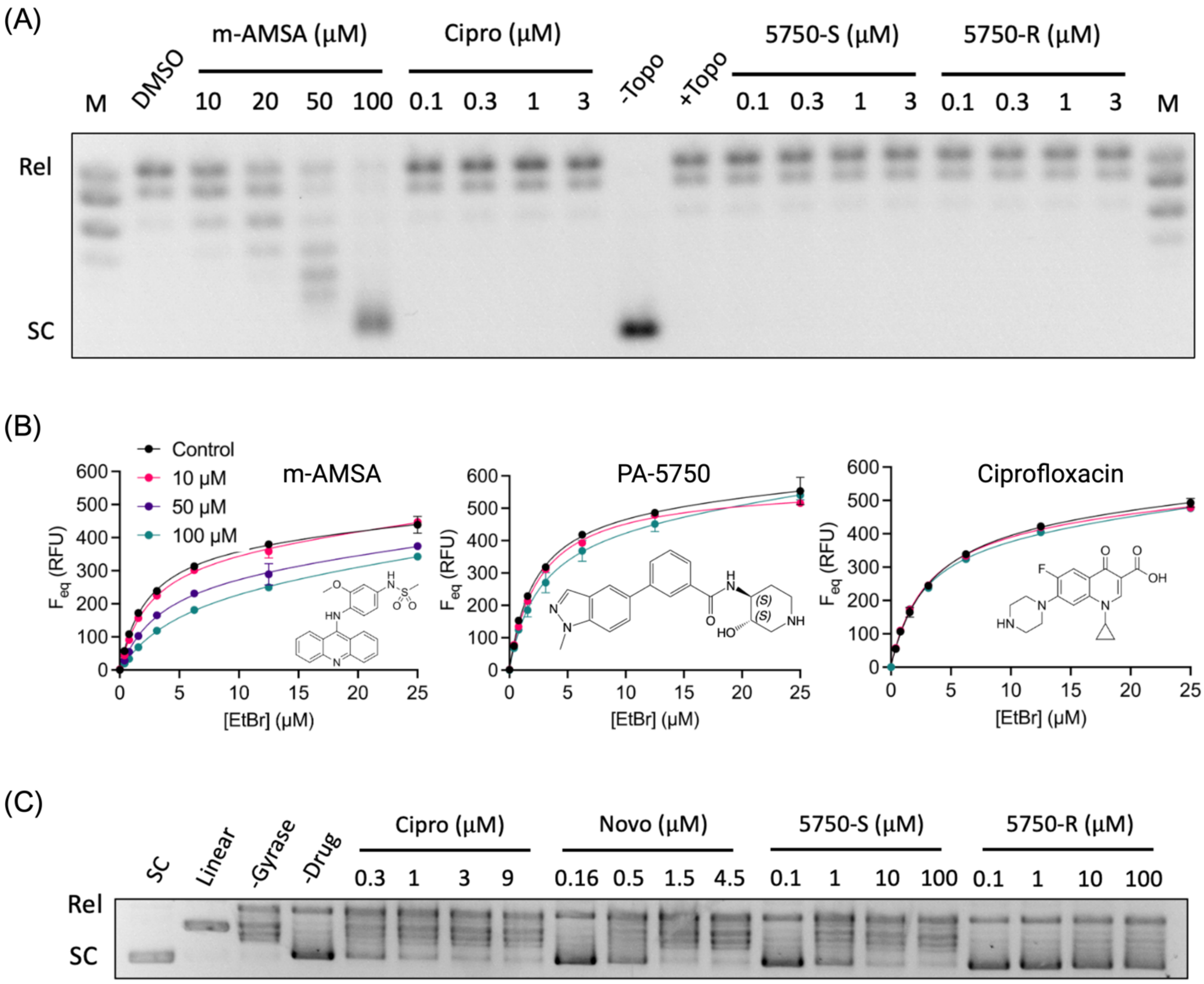


**Fig. S7.** Characterization of DNA intercalation and gyrase inhibition by PA-5750. (A) In the human topoisomerase I DNA unwinding assay, a dose-depedent shift from relaxed (Rel) to supercoiled (SC) DNA was observed for m-AMSA, a known DNA intercalator, but not for ciprofloxacin or the two enantomers of PA-5750. (B) *In vitro* ethidium bromide displacement assay showed a dose-depedent displacment of ethidium bromide from plasmid DNA with m-AMSA treatment, but no significant change after treatment with PA-5750 or ciprofloxacin. (C) In the gyrase inhibition assay, ciprofloxacin and novobiocin, both known gyrase inhibitors, exhibited dose-dependent decreases in supercoiled (SC) DNA. PA-5750 also demonstrates a dose-dependent decrease in supercoiled DNA, which is not observed for the inactive R-enantiomer.

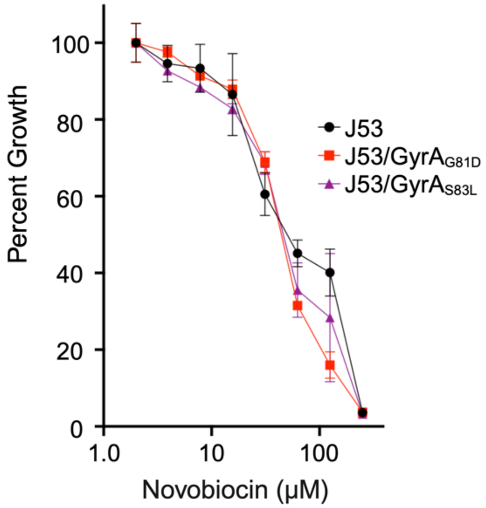


**Fig. S8.** Novobiocin MIC curves of ciprofloxacin-resistant E. coli strains. Two ciprofloxacin-resistant *E. coli* J53 mutant strains, each containing single point mutations (G81D or S83L) in GyrA, remain susceptible to the GyrB inhibitor novobiocin.

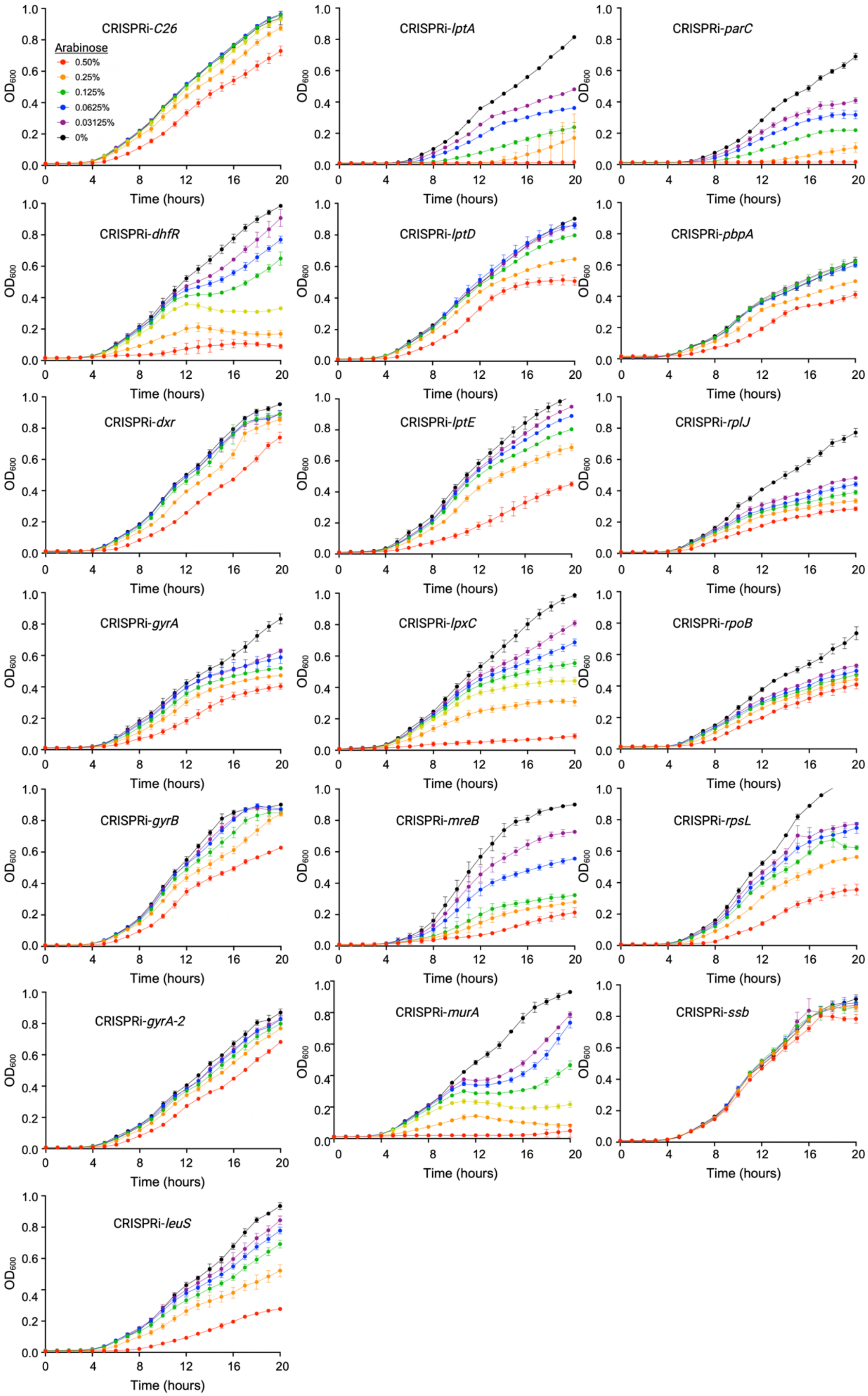
CRISPRi-C26
Arabinose
0.50%
0.25%
0.125%
0.0625%
0.03125%
0%
CRISPRi-lptA
CRISPRi-parC
CRISPRi-dhfR
CRISPRi-lptD
CRISPRi-pbpA
CRISPRi-dxr
CRISPRi-lptE
CRISPRi-rplJ
CRISPRi-gyrA
CRISPRi-lpxC
CRISPRi-rpoB
CRISPRi-gyrB
CRISPRi-mreB
CRISPRi-rpsL
CRISPRi-gyrA-2
CRISPRi-murA
CRISPRi-ssb
CRISPRi-leuS
OD600
Time (hours)

**Fig. S9.** Growth curves of CRISPRi strains. Arabinose-dose response growth curves of the 19 CRISPRi strains included in this study, including the control strain (CRISPRi-C26). In all experiments, error bars represent S.E.M of three biological replicates (n=3).

### Supplemental Tables

**Table S1:** Summary of the 39 antimicrobial compounds considered for the reference set. Antimicrobial compounds used in gene expression time trial are indicated in bold. Antibiotics exhibiting greater or equal to 4-fold shift in MIC at high inoculum are indicated by an asterisk.

| MOA | Antibiotic | Class | Target | MIC (μM) Low Inoculum | MIC (μM) High Inoculum |
|---|---|---|---|---|---|
| DNA Synthesis | **Ciprofloxacin** | **Fluoroquinolone** | **GyrA/ParC** | **0.25** | **0.25** |
| | **Levofloxacin** | **Fluoroquinolone** | **GyrA/ParC** | **1.5** | **1.5** |
| | **Novobiocin** | **Novobiocin** | **GyrB** | **>800** | **>800** |
| | **Trimethoprim** | **Anti-metabolite** | **DhfR** | **100** | **200** |
| | Cisplatin | Alkylator | DNA | 100 | 100 |
| | Hydroxyurea | Anti-metabolite | Ribonucleotide reductase | >3200 | >3200 |
| | Doxorubicin | Anthracyclines | DNA | >800 | >800 |
| | Radicicol | Macrocycle | Histidine Kinase | >400 | >400 |
| | Nitrofurantoin | Nitrofuran | DNA | >800 | >800 |
| Cell Wall Synthesis | **Meropenem*** | **Carbapenem** | **PBPs** | **1.5** | **6.25** |
| | **Imipenem** | **Carbapenem** | **PBPs** | **1.5** | **3.0** |
| | **Ceftazidime*** | **Cephalosporin** | **PBPs** | **12.5** | **400** |
| | **Piperacillin*** | **Beta Lactam** | **PBPs** | **6.25** | **>400** |
| | **Carbenicillin*** | **Beta Lactam** | **PBPs** | **200** | **3200** |
| | **Aztreonam*** | **Monobactam** | **PBPs** | **6.25** | **>400** |
| | **Fosfomycin*** | **PEP Synthesis** | **MurA** | **250** | **1600** |
| | A22* | Actin homolog | MreB | 12.5 | 400 |
| | CBR-4830* | Actin homolog | MreB | 15.63 | 62.5 |
| Protein Synthesis | **Tetracycline** | **Tetracycline** | **30S Ribosome** | **50** | **50** |
| | **Minocycline** | **Tetracycline** | **30S Ribosome** | **12.5** | **12.5** |
| | **Erythromycin** | **Macrolide** | **50S Ribosome** | **100** | **100** |
| | **Clarithromycin** | **Macrolide** | **50S Ribosome** | **100** | **100** |
| | **Amikacin** | **Aminoglycoside** | **30S Ribosome** | **1.5** | **3** |
| | **Gentamicin** | **Aminoglycoside** | **30S Ribosome** | **3** | **3** |
| | Doxycycline | Tetracycline | 30S Ribosome | 50 | 50 |
| | Tobramycin* | Aminoglycoside | 30S Ribosome | 1.56 | 6.25 |
| | Rifampicin | Ansamycins | RpoB | >400 | >400 |
| | Fusidic acid | Fusidane | FusA1 (EF-G) | >400 | >400 |
| Membrane Integrity | **Colistin*** | **Polymyxin** | **LPS** | **0.78** | **6.25** |
| | **Protegrin-1*** | **AMP** | **LPS** | **2** | **8** |
| | **PF-5081090** | **LPS Synthesis** | **LpxC** | **0.4** | **0.8** |
| | **PF-04753299** | **LPS Synthesis** | **LpxC** | **0.4** | **0.8** |
| | LL-37* | AMP | Membrane | 12.5 | >50 |
| | Fosmidomycin* | Isoprenoid biosynthesis | Dxr | 12.5 | >400 |
| | POL7001* | LPS Transport | LptD | 0.0625 | 0.500 |
| | POL7080* | LPS Transport | LptD | 0.0625 | 0.125 |
| | CHIR-090* | LPS Synthesis | LpxC | 0.78 | 3.125 |
| | MRL-494 | OMP folding | BamA | 25 | 25 |
| | Thiolactomycin | Fatty Acid Synthesis | KasA | >400 | >400 |

**Table S2:** Top 25 upregulated and downregulated annotated genes in PA14 in response to treatment with aminoglycosides, fluoroquinolones and polymyxins. Previously reported marker genes are highlighted in bold [13-20].

| Aminoglycosides | | | | | |
|---|---|---|---|---|---|
| | Upregulated | | | Downregulated | |
| Gene Locus ID | Gene Name | Z-score | Gene Locus ID | Gene Name | Z-score |
| **PA14_10420** | **tyrS** | **7.617** | **PA14_57330** | **murC** | **-8.6786** |
| **PA14_61840** | **vapI** | **7.5757** | PA14_57290 | ftsA | -8.6054 |
| PA14_62330 | phuS | 7.3815 | PA14_08760 | rpoB | -7.9094 |
| PA14_09540 | mexG | 7.2805 | PA14_57300 | ftsQ | -7.7995 |
| PA14_05840 | gcdH | 6.9018 | PA14_30660 | uvrC | -7.6364 |
| PA14_15480 | merR | 6.8436 | **PA14_29850** | **nuoN** | **-6.9983** |
| PA14_47110 | yybH | 6.5468 | PA14_23440 | orfL | -6.9223 |
| PA14_17530 | recA | 6.5229 | PA14_12100 | dacC/dacA | -6.8561 |
| PA14_29110 | cysK | 6.2604 | PA14_65350 | mutL | -6.7426 |
| PA14_65310 | hfq | 6.2598 | PA14_57320 | ddl | -6.6952 |
| **PA14_30410** | **yccA** | **6.0343** | **PA14_29880** | **nuoL** | **-6.6709** |
| **PA14_68700** | **amgR** | **5.9497** | **PA14_29860** | **nuoM** | **-6.6183** |
| PA14_72560 | np20 | 5.8613 | PA14_30310 | lolA | -6.5492 |
| PA14_47430 | czcD | 5.7271 | **PA14_57340** | **murG** | **-6.5487** |
| PA14_08340 | trpG | 5.719 | PA14_66200 | wapQ | -6.5148 |
| PA14_66330 | msrA | 5.6624 | PA14_57360 | ftsW | -6.1145 |
| PA14_02550 | mdcA | 5.6271 | PA14_61740 | lolB | -6.0926 |
| PA14_69670 | lysA | 5.5666 | PA14_41160 | nppD | -6.0837 |
| PA14_65300 | hflX | 5.5591 | PA14_23450 | orfM | -6.004 |
| PA14_08480 | argC | 5.5048 | PA14_57275 | ftsZ | -5.8326 |
| PA14_72200 | prfH | 5.4321 | **PA14_57370** | **murD** | **-5.8243** |
| PA14_27020 | mttC | 5.3618 | PA14_61700 | prfA | -5.7578 |
| PA14_51050 | pcd | 5.3079 | PA14_70570 | recG | -5.7172 |
| PA14_04390 | ygdP | 5.2917 | **PA14_29900** | **nuoJ** | **-5.6437** |
| PA14_16720 | pemA | 5.1794 | **PA14_29920** | **nuoI** | **-5.6178** |
| **Fluoroquinolones** | | | | | |
| | Upregulated | | | Downregulated | |
| Gene Locus ID | Gene Name | Z-score | Gene Locus ID | Gene Name | Z-score |
| PA14_08010 | gpV | 10 | **PA14_51880** | **oprD** | **-5.3404** |
| PA14_08020 | gpW | 10 | PA14_08760 | rpoB | -5.2747 |
| PA14_08040 | gpI | 10 | **PA14_57275** | **ftsZ** | **-4.922** |
| PA14_08070 | gpFI | 10 | PA14_57330 | murC | -4.7292 |
| PA14_08090 | gpFII | 10 | PA14_08820 | fusA1 | -4.5735 |
| PA14_08130 | gpU | 10 | **PA14_23470** | **wbpM** | **-4.4856** |
| PA14_08140 | XR2 | 10 | PA14_09050 | secY | -4.4069 |
| PA14_08150 | gpD | 10 | **PA14_56070** | **mvaT** | **-4.3601** |
| PA14_08160 | lys | 10 | **PA14_57290** | **ftsA** | **-4.0381** |
| PA14_08300 | JF1 | 10 | PA14_57320 | ddl | -3.8933 |
| **PA14_17530** | **recA** | **10** | **PA14_57300** | **ftsQ** | **-3.8865** |
| **PA14_17540** | **recX** | **10** | PA14_44670 | zipA | -3.5393 |
| PA14_49520 | pyoS3A | 10 | **PA14_41210** | **hupB** | **-3.5233** |
| **PA14_07950** | **prtN** | **9.6405** | PA14_58390 | dppA3 | -3.455 |
| **PA14_19950** | **yebG** | **8.8869** | PA14_08680 | tufB | -3.3908 |
| **PA14_07960** | **prtR** | **8.0788** | PA14_08540 | mepM | -3.1602 |
| PA14_55610 | dnaE2 | 7.9149 | PA14_41570 | oprF | -3.0695 |
| PA14_49510 | pyoS3I | 7.4688 | PA14_09900 | prpL | -3.0404 |
| PA14_53820 | ampDh3 | 6.7152 | PA14_23440 | orfL | -2.9981 |
| PA14_08360 | trpC | 5.7091 | PA14_57450 | mraW | -2.9755 |
| **PA14_63010** | **recN** | **5.4647** | PA14_57340 | murG | -2.9626 |
| **PA14_25160** | **lexA** | **4.9485** | PA14_23370 | orfK | -2.9013 |
| PA14_08340 | trpG | 4.8995 | PA14_66670 | ponA | -2.8365 |
| PA14_08350 | trpD | 4.8875 | PA14_23450 | orfM | -2.8156 |
| PA14_14590 | queA | 4.3626 | PA14_57370 | murD | -2.7992 |

| **Polymyxins** | | | | | |
|---|---|---|---|---|---|
| Upregulated | | | Downregulated | | |
| Gene Locus ID | Gene Name | Z-score | Gene Locus ID | Gene Name | Z-score |
| **PA14_09520** | **mexI** | **9.6027** | PA14_10540 | fixG | -9.9006 |
| **PA14_09540** | **mexG** | **9.5397** | PA14_06720 | **nirF** | **-9.6825** |
| PA14_09500 | opmD | 9.5387 | **PA14_20200** | **nosZ** | **-9.4315** |
| **PA14_09270** | **pchE** | **7.2334** | **PA14_06750** | **nirS** | **-9.4255** |
| PA14_35160 | dsbA2 | 7.0763 | **PA14_06840** | **norD** | **-9.0603** |
| **PA14_60820** | **oprJ** | **6.4143** | PA14_10550 | cysI | -8.8956 |
| PA14_25080 | fadB | 6.2748 | **PA14_06810** | **norC** | **-8.6183** |
| PA14_62400 | yfdZ | 5.8538 | **PA14_20230** | **nosR** | **-8.5103** |
| **PA14_63160** | **pmrB** | **5.7799** | **PA14_06670** | **nirJ** | **-7.0148** |
| PA14_67090 | mdoG | 5.7571 | **PA14_06710** | **nirD** | **-6.515** |
| PA14_21370 | fadD1 | 5.4661 | PA14_52040 | purM | -6.3151 |
| PA14_25090 | fadA | 5.3336 | PA14_57590 | rplM | -6.2624 |
| PA14_27520 | gpo | 4.841 | **PA14_06700** | **nirL** | **-5.9319** |
| PA14_62330 | phuS | 4.7729 | **PA14_06650** | **nirN** | **-5.9222** |
| **PA14_18360** | **arnC** | **4.7144** | **PA14_06680** | **nirH** | **-5.8314** |
| PA14_05510 | yceI | 4.714 | PA14_68350 | arcC | -5.8095 |
| PA14_39150 | accC | 4.6154 | PA14_10500 | ccoN | -5.4014 |
| **PA14_05530** | **mexA** | **4.5963** | **PA14_06690** | **nirG** | **-5.3174** |
| **PA14_63150** | **pmrA** | **4.5841** | PA14_62780 | yhbC | -5.0473 |
| **PA14_09530** | **mexH** | **4.5637** | **PA14_06660** | **nirE** | **-4.9944** |
| PA14_05520 | mexR | 4.5546 | PA14_72460 | cc4 | -4.836 |
| **PA14_49180** | **phoP** | **4.3012** | PA14_04790 | laoB | -4.7811 |
| PA14_68700 | ompR | 4.2781 | **PA14_06790** | **nirO** | **-4.7043** |
| PA14_29400 | rhs | 4.1427 | PA14_04810 | laoC | -4.4951 |
| PA14_68900 | fbpA | 4.12 | PA14_15990 | trmD | -4.4671 |

**Table S3:** Summary of growth phenotypes and antimicrobial susceptibility of hypomorphic strains. Constitutive promoters are ordered from weakest to strongest, with $P_{Pa1}$ being weakest and $P_{Pa5}$ being the strongest, as described previously [1].

| Mutant/Strain | Promoter (Location) | Gene Function | Gene Expression[a] $log_2$FoldChange ($p_{adj}$) | Treatments | Classification | MIC (μM) Low Inoculum | MIC (μM) High Inoculum |
|---|---|---|---|---|---|---|---|
| PA14 | N/A | NA | NA | POL7080 | NA | 0.0625 | 0.125 |
| | | | | Doxycycline | NA | 50 | 50 |
| | | | | Ciprofloxacin | NA | 0.25 | 0.25 |
| | | | | MRL-494 | NA | 62.5 | >50 |
| | | | | Fosfomycin | NA | 250 | 1600 |
| | | | | PF-04753299 | NA | 0.4 | 0.8 |
| | | | | Trimethoprim | NA | 100 | 200 |
| | | | | Fosmidomycin | NA | 12.5 | >500 |
| *lptD*-hypomorph | $P_{araBAD}$::*lptD-surA-pdxA* (native site) | LPS Transport | -4.08 (< $2.0x10^{-308}$) | POL7080 | On-target | < 0.004 | 0.03125 |
| | | | | Doxycycline | Off-target | < 3.125 | 25 |
| | | | | Ciprofloxacin | Off-target | 0.125 | 0.125 |
| *dxr*-hypomorph | $P_{araBAD}$::*dxr* (*attB-Tn7* site) | Isoprenoid Synthesis | -4.50 ($1.0x10^{-238}$) | Fosmidomycin | On-target | 12.5 | >400 |
| | | | | Doxycycline | Off-target | 25 | 50 |
| | | | | Ciprofloxacin | Off-target | 0.5 | 0.25 |
| *murA*-hypomorph | $P_{Pa3}$::*murA* (*attB-Tn7* site) | Peptidoglycan Synthesis | -1.69 ($2.6 x10^{-90}$) | Fosfomycin | On-target | 16 | 400 |
| | | | | Doxycycline | Off-target | 50 | 50 |
| | | | | Ciprofloxacin | Off-target | 0.25 | 0.5 |
| *gyrA*-hypomorph | $P_{Pa1}$:*gyrA* (*attB-Tn7* site) | DNA Gyrase | 0.3 ($4.0x10^{-7}$) | Ciprofloxacin | On-target | 0.25 | 0.25 |
| | | | | Doxycycline | Off-target | 25 | 25 |
| | | | | PF-04753299 | Off-target | 0.5 | 0.5 |
| *lpxC*-hypomorph | $P_{Pa4}$:*lpxC* (*attB-Tn7* site) | LPS Synthesis | -3.2 ($9.4x10^{-251}$) | PF-04753299 | On-target | 0.5 | 0.5 |
| | | | | Doxycycline | Off-target | 12.5 | 25 |
| | | | | Ciprofloxacin | Off-target | 0.5 | 0.25 |
| *dhfR*-hypomorph | $P_{Pa5}$::*dhfR* (*attB-Tn7* site) | Folate Metabolism | 3.5 (NA[b]) | Trimethoprim | On-target | 200 | 200 |
| | | | | Doxycycline | Off-target | 25 | 50 |
| | | | | POL7080 | Off-target | 0.125 | 0.125 |
| *bamA*-hypomorph | $P_{Pa3}$:*bamA* (*attB-Tn7* site) | OMP Assembly | -0.9 ($8.8x10^{-45}$) | MRL-494 | On-target | 3.125 | 12.5 |
| | | | | Doxycycline | Off-target | 25 | 25 |
| | | | | Ciprofloxacin | Off-target | 0.125 | 0.125 |

[a]DESeq2 relative to gene expression to PA14 without inducer with adjusted p-values shown in parentheses. | [b]Filtered out by DESeq2 for having a low mean normalized count

**Table S4:** Target and MOA prediction of antimicrobial compounds from external and internal datasets using the PerSpecTM approach.

| **Compound** | **Predicted Target** | **Predicted MOA** | $r_{predict}$ | **Target / MOA PPV** | **Closest Reference** |
|---|---|---|---|---|---|
| External Validation using Murray et al. (2015) **[21]** | | | | | |
| Peroxide | GyrA/ParC | DNA Synthesis | 0.51 | 81% / 96% | Ciprofloxacin |
| Gentamicin | 30S RS | Protein Synthesis | 0.48 | 79% / 96% | Tobramycin |
| Ciprofloxacin | GyrA/ParC | DNA Synthesis | 0.33 | 61% / 94% | Levofloxacin |
| Neomycin | 30S RS | Protein Synthesis | 0.33 | 61% / 94% | Amikacin |
| Bleach | DNA | DNA Synthesis | 0.26 | NA / NA | Nitrofurantoin |
| Polymyxin B | LptD | Membrane Integrity | 0.26 | NA / NA | POL7080 |
| Target and MOA predictions of internal screening hits | | | | | |
| PA-0918 | LPS | Membrane Integrity | 0.74 | 100% / 100% | Colistin |
| PA-69180 | MreB | Cell wall Synthesis | 0.38 | 68% / 94% | A22 |
| PA-5750 | GyrA/ParC | DNA Synthesis | 0.56 | 88% / 96% | Ciprofloxacin |

**Table S5:** Chemical structures and MIC data for screening hits PA-0918, PA-69180 and PA-5750. The fold decrease in MIC relative to PA14 is shown in parentheses.

| Screening Hit | Screening Strain | Chemical Structure | PA14 | MIC (μM) Screening Strain | PAO397 |
|---|---|---|---|---|---|
| PA-0918 | *lptD*-hypomorph | 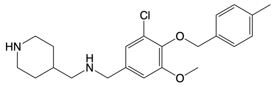 | 250 | 125 (2) | 31.2 (8) |
| PA-69180 | *oprL*-hypomorph | 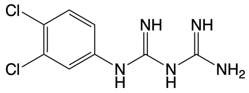 | >250 | 31.2 (>8) | 15.6 (>16) |
| PA-5750 | *lptE*-hypomorph | 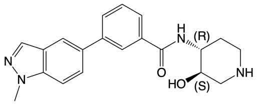 | 250 | 125 (2) | <15.6 (>16) |

**Table S6:** Summary of resistant mutants sequenced after selection with PA-69180. The fold change in MIC relative to PAO397 is shown in parentheses.

| Strain | PAO1_ID | Gene | SNPs | Protein Change | Function | MIC in LB (μM) | | |
|---|---|---|---|---|---|---|---|---|
| | | | | | | PA-69180 | A22 | Ciprofloxacin |
| PAO397 | - | - | - | - | - | 15.6 | 2.34 | 0.0156 |
| PAO397-r1 | PA4481 | *mreB* | C337T | P113S | Actin Homolog | 500 (32) | 150 (64) | 0.0156 (1) |
| | PA1874 | *hypo*[a] | delC4830-G5074 | A1609[Trunc‡] | Putative O-acetylesterase | | | |
| PAO397-r2 | PA4481 | *mreB* | C489A | D163E | Actin Homolog | 500 (32) | 150 (64) | 0.0156 (1) |

[a]Hypothetical protein | Trunc‡: frameshift after amino acid causing mistranslation with premature truncation

**Table S7**: Summary of CRISPRi strains used in this study, including guide sequence information and expressional knockdown levels.

| Strain | Gene Function | PAM[a] | Guide Sequence | Location | Arabinose (%) | 360-minute Gene Expression[b] $log_2$FoldChange ($p_{adj}$) |
|---|---|---|---|---|---|---|
| *CRISPRi-C26* | NA | NA | GTATAACACATATTAGTGTAT | Negative control guide | 0% | NA |
| | | | | | 0.125% | NA |
| | | | | | 0.5% | NA |
| *CRISPRi-dhfR* | Folate Metabolism | NNGCGA | ACCTCATCGTGGAAGGACAAAG | 28bp upstream of *dhfR* | 0% | 0.71 (1.0 x $10^{0}$) |
| | | | | | 0.125% | -7.44 (4.5 x $10^{-2}$) |
| | | | | | 0.5% | -3.28 (5.6 x $10^{-1}$) |
| *CRISPRi-dxr* | Isoprenoid Synthesis | NNGCGA | GCCAGGCGGCTGAAGCCGGTCA | 106bp into *dxr* | 0% | -1.49 (1.7 x $10^{-1}$) |
| | | | | | 0.125% | -1.61 (5.5 x $10^{-2}$) |
| | | | | | 0.5% | -1.69 (1.7 x $10^{-2}$) |
| *CRISPRi-gyrA* | DNA Gyrase | NNGCGA | GGATGTTGGTCGCCATGCCCA | 529bp into *gyrA* | 0% | -0.071 (9.7 x $10^{-1}$) |
| | | | | | 0.125% | -1.048 (8.6 x $10^{-6}$) |
| | | | | | 0.5% | -0.62 (2.2 x $10^{-2}$) |
| *CRISPRi-gyrA-2* | DNA Gyrase | NNGTGA | GCTTGTTCAACTGGTACGGCAGCT | 801bp into *gyrA* | 0% | -0.04 (1.0 x $10^{0}$) |
| | | | | | 0.125% | -0.08 (1.0 x $10^{0}$ |
| | | | | | 0.5% | 0.27 (9.3 x $10^{-1}$) |
| *CRISPRi-gyrB* | DNA Gyrase | NNGTGA | GTGATCGACTCATCCGTATGGA | 194bp into *gyrB* | 0% | -1.44 (7.6 x $10^{-9}$) |
| | | | | | 0.125% | -1.40 (1.3 x $10^{-7}$) |
| | | | | | 0.5% | -1.68 (2.9 x $10^{-12}$) |
| *CRISPRi-leuS* | Leucyl-tRNA synthetase | NNGTGA | GGTAGTAGTCCGGCTTGCAGG | 380bp into *leuS* | 0% | -0.65 (1.0 x $10^{0}$) |
| | | | | | 0.125% | -1.59 (3.9 x $10^{-5}$) |
| | | | | | 0.5% | -2.30 (2.8 x $10^{-13}$) |
| *CRISPRi-lptA* | LPS Transport | NNGCGA | ACCACGACGTCGCCGCGGTAGA | 149bp into *lptA* | 0% | -1.36 (6.9 x $10^{-2}$) |
| | | | | | 0.125% | -1.64 (2.5 x $10^{-4}$) |
| | | | | | 0.5% | -1.92 (1.9 x $10^{-5}$) |
| *CRISPRi-lptD* | LPS Transport | NNGTGA | GCTGCAGAGCCAGCAGGCTGC | 52bp into *lptD* | 0% | -2.02 (5.3 x $10^{-14}$) |
| | | | | | 0.125% | -3.56 (3.0 x $10^{-37}$) |
| | | | | | 0.5% | -3.27 (3.4 x $10^{-35}$) |
| *CRISPRi-lptE* | LPS Transport | NNGTGA | ACCAGGTGGTAGGGCGCGTTGC | 194bp into *lptE* | 0% | -0.08 (1.0 x $10^{0}$) |
| | | | | | 0.125% | -0.65 (1.0 x $10^{0}$) |
| | | | | | 0.5% | -1.40 (6.3 x $10^{-1}$) |

| | | | | | | |
|---|---|---|---|---|---|---|
| *CRISPRi-lpxC* | LPS Synthesis | NNGTAA | ATCGGCGCCGGTTTCAGGGTC | 77bp into *lpxC* | 0% | -2.65 (1.7 x $10^{-21}$) |
| | | | | | 0.125% | -4.43 (2.5 x $10^{-62}$) |
| | | | | | 0.5% | -4.87 (3.0 x $10^{-73}$) |
| *CRISPRi-mreB* | Rod-shape Morphology | NNGCGA | GACGCTCTTCTGGCTGCCATGG | 121bp into *mreB* | 0% | -2.14 (3.0 x $10^{-5}$) |
| | | | | | 0.125% | -2.95 (4.5 x $10^{-11}$) |
| | | | | | 0.5% | -3.53 (4.3 x $10^{-15}$) |
| *CRISPRi-murA* | Peptidoglycan Synthesis | NNGTGA | GCAGGTGCGGCAGGTTGCAGA | 121bp into *murA* | 0% | -2.23 (1.4 x $10^{-17}$) |
| | | | | | 0.125% | -2.61 (3.9 x $10^{-24}$) |
| | | | | | 0.5% | -3.26 (6.8 x $10^{-36}$) |
| *CRISPRi-parC* | DNA Topoisomerase | NNGTGA | AATAATTCAGATAGGCCTGCT | 65bp into *parC* | 0% | -2.41 (1.5 x $10^{-7}$) |
| | | | | | 0.125% | -2.49 (2.1 x $10^{-7}$) |
| | | | | | 0.5% | -2.55 (9.6 x $10^{-6}$) |
| *CRISPRi-pbpA* | Peptidoglycan Synthesis | NNGTGA | GCAGGTCCTCGGTGCGTTCGC | 275bp into *pbpA* | 0% | -1.78 (2.4 x $10^{-3}$) |
| | | | | | 0.125% | -1.07 (1.5 x $10^{-1}$) |
| | | | | | 0.5% | -1.74 (8.2 x $10^{-3}$) |
| *CRISPRi-rplJ* | 50S Ribosomal Subunit | NNGCGA | GCTTTGGCAGCCTCGTTGACTT | 41bp into *rplJ* | 0% | 0.00 (1.0 x $10^{0}$) |
| | | | | | 0.125% | -0.58 (1.4 x $10^{-2}$) |
| | | | | | 0.5% | -0.66 (1.9 x $10^{-3}$) |
| *CRISPRi-rpoB* | RNA Polymerase | NNGTGA | GTTCCAGCTGGTTGATGTGGCG GG | 827bp into *rpoB* | 0% | -0.67 (3.7 x $10^{-3}$) |
| | | | | | 0.125% | -0.23 (7.2 x $10^{-1}$) |
| | | | | | 0.5% | 0.25 (3.9 x $10^{-1}$) |
| *CRISPRi-rpoB* | 30S Ribosomal Subunit | NNGTGA | ACTACGCTGTGCTCTTGCAGG | 219bp into *rpsL* | 0% | -0.54 (4.0 x $10^{-1}$) |
| | | | | | 0.125% | -1.05 (2.2 x $10^{-5}$) |
| | | | | | 0.5% | -1.55 (3.6 x $10^{-12}$) |
| *CRISPRi-ssb* | DNA Replication and Repair | NNGTGA | GCTTCCAGCTCTCGCTGGTGGC GA | 101bp into *ssb* | 0% | 0.07 (1.0 x $10^{0}$) |
| | | | | | 0.125% | 0.24 (1.0 x $10^{0}$) |
| | | | | | 0.5% | 0.91 (1.5 x $10^{-1}$) |

[a]Protospacer adjacent motif | [b]DESeq2 relative to CRISPRi-*C26* control strain at matched arabinose dose with adjusted p-values shown in parentheses.

| **Table S8**: Strains used in this study. $P_{araBAD}$ indicates the *araBAD* operon. Constitutive promoters are ordered from weakest to strongest, with $P_{Pa1}$ being weakest and $P_{Pa5}$ being the strongest [1]. | | |
|---|---|---|
| Strain | Relevant Genotype | Source |
| *PA14* | Wildtype *P. aeruginosa* strain | Hung Lab [1] |
| *lptD*-hypomorph | *PA14, $P_{araBAD}$::lptD-surA-pdxA* | This study |
| *dxr*-hypomorph | *PA14, $P_{araBAD}$::dxr(attB-Tn7) Δdxr* [a] | This study |
| *murA*-hypomorph | *PA14, $P_{Pa3}$::murA* | Hung Lab [1] |
| *gyrA*-hypomorph | *PA14, $P_{Pa1}$::gyrA* | Hung Lab [1] |
| *lpxC*-hypomorph | *PA14, $P_{Pa4}$::lpxC* | Hung Lab [1] |
| *dhfR*-hypomorph | *PA14, $P_{Pa5}$::dhfR* | Hung Lab [1] |
| *bamA*-hypomorph | *PA14, $P_{Pa3}$::bamA* | Hung Lab [1] |
| *lptE*-hypomorph | *PA14, $P_{araBAD}$::lptE(attB-Tn7) ΔlptE* [a] | This study |
| *oprL*-hypomorph | *PA14, $P_{araBAD}$::oprL(attB-Tn7) ΔoprL* [a] | This study |
| *CRISPRi-C26* | *PA14, $P_{araBAD}$::dCas9$_{spas}$(attB-Tn7)* | This study |
| *CRISPRi-dhfR* | *PA14, $P_{araBAD}$::dCas9$_{spas}$(attB-Tn7)* | This study |
| *CRISPRi-dxr* | *PA14, $P_{araBAD}$::dCas9$_{spas}$(attB-Tn7)* | This study |
| *CRISPRi-gyrA* | *PA14, $P_{araBAD}$::dCas9$_{spas}$(attB-Tn7)* | This study |
| *CRISPRi-gyrA-2* | *PA14, $P_{araBAD}$::dCas9$_{spas}$(attB-Tn7)* | This study |
| *CRISPRi-gyrB* | *PA14, $P_{araBAD}$::dCas9$_{spas}$(attB-Tn7)* | This study |
| *CRISPRi-leuS* | *PA14, $P_{araBAD}$::dCas9$_{spas}$(attB-Tn7)* | This study |
| *CRISPRi-lptA* | *PA14, $P_{araBAD}$::dCas9$_{spas}$(attB-Tn7)* | This study |
| *CRISPRi-lptD* | *PA14, $P_{araBAD}$::dCas9$_{spas}$(attB-Tn7)* | This study |
| *CRISPRi-lptE* | *PA14, $P_{araBAD}$::dCas9$_{spas}$(attB-Tn7)* | This study |
| *CRISPRi-lpxC* | *PA14, $P_{araBAD}$::dCas9$_{spas}$(attB-Tn7)* | This study |
| *CRISPRi-mreB* | *PA14, $P_{araBAD}$::dCas9$_{spas}$(attB-Tn7)* | This study |
| *CRISPRi-murA* | *PA14, $P_{araBAD}$::dCas9$_{spas}$(attB-Tn7)* | This study |
| *CRISPRi-parC* | *PA14, $P_{araBAD}$::dCas9$_{spas}$(attB-Tn7)* | This study |
| *CRISPRi-pbpA* | *PA14, $P_{araBAD}$::dCas9$_{spas}$(attB-Tn7)* | This study |
| *CRISPRi-rplJ* | *PA14, $P_{araBAD}$::dCas9$_{spas}$(attB-Tn7)* | This study |
| *CRISPRi-rpoB* | *PA14, $P_{araBAD}$::dCas9$_{spas}$(attB-Tn7)* | This study |
| *CRISPRi-rpsL* | *PA14, $P_{araBAD}$::dCas9$_{spas}$(attB-Tn7)* | This study |
| *CRISPRi-ssb* | *PA14, $P_{araBAD}$::dCas9$_{spas}$(attB-Tn7)* | This study |
| PAO397 | PAO1, Δ(*mexEF-oprN*)[b] Δ*opmH*362[b] Δ(*mexAB-oprM*)[b] *nfxB* Δ(*mexCD-oprJ*)[b] Δ(*mexJKL*)[b] Δ(*mexXY*)[b] OpmH+ | Schweizer Lab [22] |
| MG1655 | *rph-1 ilvG rfb-50* | Low Lab [23] |
| MG1655/FtsZup | *MG1655+pTB63, $P_{native}$::ftsQAZ(episomal)* | Bernhardt Lab [10] |
| J53 | J53 encoding wildtype GyrA | Hung Lab [24] |
| J53/$GyrA_{G81D}$ | J53 encoding GyrA with G81D mutation | Hung Lab [24] |
| J53/$GyrA_{S83L}$ | J53 encoding GyrA with S83L mutation | Hung Lab [24] |
| [a]Native gene deleted but replaced by KanR cassette \| [b]Unmarked deletion but containing *FRT* scar | | |

| Table S9: Plasmids used in this study. | | | |
|---|---|---|---|
| Plasmid | Relevant Properties[a] | Ori[b] | Source |
| *pRK2013* | KanR; plasmid mobilization helper | ColE1 | Helinski Lab [25] |
| *pTNS3* | AmpR; *tnsABCD* from *P1* and $P_{lac}$ | R6Kγ | Schweizer Lab [26] |
| *pEXG2-AmpR-dxr* | AmpR; $P_{sacB}$:*sacB* \|$dxr^{up}$:*KanR*:$dxr^{Down}$ | ColE1 | This study |
| *pEXG2-AmpR-lptD* | AmpR; $P_{sacB}$:*sacB* \|*Pc::KanR*-$P_{araBAD}$::*lptD-surA-pxdA* | ColE1 | This study |
| *pEXG2-AmpR-oprL* | AmpR; $P_{sacB}$:*sacB* \| $oprL^{up}$:*KanR*:$oprL^{Down}$ | ColE1 | This study |
| *pEXG2-AmpR-lptE* | AmpR; $P_{sacB}$:*sacB* \|$P_{araBAD}$: $lptE^{up}$:*KanR*:$lptE^{Down}$ | ColE1 | This study |
| *pUC18-mini-Tn7-GmR-oprL* | *GmR*; $P_{araBAD}$::*oprL* | pMB1 | Hung Lab [1] |
| *pUC18-mini-Tn7-GmR-dxr* | *GmR*; $P_{araBAD}$::*dxr* | pMB1 | This study |
| *pUC18-mini-Tn7-GmR-lptE* | *GmR*; $P_{araBAD}$::*lptE* | pMB1 | This study |
| *pUC18-mini-Tn7-GmR-CRISPRI-C26* | *GmR*; $P_{araBAD}$::$dCas9_{spas}$ \| $P_{ProD}$::$sgRNA_{(C26)}$ | pMB1 | This study |
| *pUC18-mini-Tn7-GmR-CRISPRi-dhfR* | *GmR*; $P_{araBAD}$::$dCas9_{spas}$ \| $P_{ProD}$::$sgRNA_{(dhfR)}$ | pMB1 | This study |
| *pUC18-mini-Tn7-GmR-CRISPRi-dxr* | *GmR*; $P_{araBAD}$::$dCas9_{spas}$ \| $P_{ProD}$::$sgRNA_{(dxr)}$ | pMB1 | This study |
| *pUC18-mini-Tn7-GmR-CRISPRi-lptD* | *GmR*; $P_{araBAD}$::$dCas9_{spas}$ \| $P_{ProD}$::$sgRNA_{(lptD)}$ | pMB1 | This study |
| *pUC18-mini-Tn7-GmR-CRISPRi-lpxC* | *GmR*; $P_{araBAD}$::$dCas9_{spas}$ \| $P_{ProD}$::$sgRNA_{(lpxC)}$ | pMB1 | This study |
| *pUC18-mini-Tn7-GmR-CRISPRi-mreB* | *GmR*; $P_{araBAD}$::$dCas9_{spas}$ \| $P_{ProD}$::$sgRNA_{(mreB)}$ | pMB1 | This study |
| *pUC18-mini-Tn7-GmR-CRISPRi-murA* | *GmR*; $P_{araBAD}$::$dCas9_{spas}$ \| $P_{ProD}$::$sgRNA_{(murA)}$ | pMB1 | This study |
| *pUC18-mini-Tn7-GmR-CRISPRi-gyrA* | *GmR*; $P_{araBAD}$::$dCas9_{spas}$ \| $P_{ProD}$::$sgRNA_{(gyrA)}$ | pMB1 | This study |
| *pUC18-mini-Tn7-GmR-CRISPRi-gyrA-2* | *GmR*; $P_{araBAD}$::$dCas9_{spas}$ \| $P_{ProD}$::$sgRNA_{(gyrA-2)}$ | pMB1 | This study |
| *pUC18-mini-Tn7-GmR-CRISPRi-gyrB* | *GmR*; $P_{araBAD}$::$dCas9_{spas}$ \| $P_{ProD}$::$sgRNA_{(gyrB)}$ | pMB1 | This study |
| *pUC18-mini-Tn7-GmR-CRISPRi-parC* | *GmR*; $P_{araBAD}$::$dCas9_{spas}$ \| $P_{ProD}$::$sgRNA_{(parC)}$ | pMB1 | This study |
| *pUC18-mini-Tn7-GmR-CRISPRi-ssb* | *GmR*; $P_{araBAD}$::$dCas9_{spas}$ \| $P_{ProD}$::$sgRNA_{(ssb)}$ | pMB1 | This study |
| *pUC18-mini-Tn7-GmR-CRISPRi-lptA* | *GmR*; $P_{araBAD}$::$dCas9_{spas}$ \| $P_{ProD}$::$sgRNA_{(lptA)}$ | pMB1 | This study |
| *pUC18-mini-Tn7-GmR-CRISPRi-lptE* | *GmR*; $P_{araBAD}$::$dCas9_{spas}$ \| $P_{ProD}$::$sgRNA_{(lptE)}$ | pMB1 | This study |
| *pUC18-mini-Tn7-GmR-CRISPRi-rpsL* | *GmR*; $P_{araBAD}$::$dCas9_{spas}$ \| $P_{ProD}$::$sgRNA_{(rpsL)}$ | pMB1 | This study |
| *pUC18-mini-Tn7-GmR-CRISPRi-rplJ* | *GmR*; $P_{araBAD}$::$dCas9_{spas}$ \| $P_{ProD}$::$sgRNA_{(rplJ)}$ | pMB1 | This study |
| *pUC18-mini-Tn7-GmR-CRISPRi-rpoB* | *GmR*; $P_{araBAD}$::$dCas9_{spas}$ \| $P_{ProD}$::$sgRNA_{(rpoB)}$ | pMB1 | This study |
| *pUC18-mini-Tn7-GmR-CRISPRi-leuS* | *GmR*; $P_{araBAD}$::$dCas9_{spas}$ \| $P_{ProD}$::$sgRNA_{(leuS)}$ | pMB1 | This study |
| *pUC18-mini-Tn7-GmR-CRISPRi-pbpA* | *GmR*; $P_{araBAD}$::$dCas9_{spas}$ \| $P_{ProD}$::$sgRNA_{(pbpA)}$ | pMB1 | This study |

[a]*P1*, $P_{lac}$, $P_{araBAD}$, and $P_{ProD}$ are *E. coli P1* promoter, *lac* operon, *araBAD* operon, and *ProD* promoter, respectively. KanR, AmpR, and GmR denote kanamycin, ampicillin, and gentamicin resistance cassettes, respectively.
[b]origin of replication

**Table S10**: Gene blocks and primers used in this study. Lowercase bases are gateway sequences.

| | | |
|---|---|---|
| **Gene Blocks** | | |
| **Name** | **Sequence[a]** | |
| 1. Spas-handle | GGTTAAGCTTTTCTAGAGCACAGCTAACACCACGTCGTCCCTATCTGCTGCCCTAGGTCTATGAGTGGTTGCTGGATAACTTTACGGGCATGCATAAGGCTCGTATAATATATTCAGGGAGACCACAACGGTTTCCCTCTACAAATAATTTTGTTTAACTTTTACTAGAGTCACACAGGAAAGTACTAGGAGACGATTAATGCGTCTCGGTTTTTGTACTCGAAAGAGCCTACAAAGATAAGGCTTTATGCCGAATTCAAGCACCCCATGTTTTGACATGAGGTGCTTTTTTTTGATAAAACGAAAGGCCCAGTCTTTCGACTGAGCCTTTCGTTTTATAAACGGGGTACCTTGG | |
| **Primers** | | |
| **Name** | **Description** | **Sequence[a]** |
| 1. pEXG2-dxr-upstream-FOR | Forward primer to amplify upstream region of *dxr* with pEXG2-AMP overhang for assembly of *pEXG2-AmpR-dxr* | TGCGGTATTTCACACCGCATATGCAGGAAACAGCTATGACCGGATCATCACGGCGCTGGT |
| 2. pEXG2-dxr-upstream-REV | Reverse primer to amplify upstream region of *dxr* with kanamycin resistance cassette overhang for assembly of *pEXG2-AmpR-dxr* | GCGTGCAATCCATCTTGTTCAATCATGGCGCACCCCAGCCGGC |
| 3. pEXG2-dxr-downstream-FOR | Forward primer to amplify downstream region of *dxr* with kanamycin resistance cassette overhang for assembly of *pEXG2-AmpR-dxr* | GCGCATCGCCTTCTATCGCCTTCTTGACGAGTTCTTCTGATCTTCGGACGGCCTGGAG |
| 4. pEXG2-dxr-downstream-REV | Reverse primer to amplify downstream region of *dxr* with pEXG2-AMP overhang for assembly of *pEXG2-AmpR-dxr* | AGCATAAATGTAAAGCAAGCTTTGTAAAACGACGGCCAGTGCTGCCAGTCGTCGACG |
| 5. pEXG2-lptD-upstream-FOR | Forward primer to amplify upstream region of native promoter of *lptD* operon. Lowercase bases are gateway sequences. | tacaaaaaagcaggctTGCATGCTGCCCTGGCG |
| 6. pEXG2-lptD-upstream-REV | Reverse primer to amplify upstream region of native promoter of *lptD* operon. | GTTTAACTTTAAGAAGGAGATATACATACCCATGGCAGTGAAATCCCTCGTG |
| 7. pEXG2-lptD-downstream-FOR | Forward primer to amplify downstream region of native promoter of *lptD* operon. | CTTGGTGTATCCAACGGCGTATGTCTGATGATGCCCGTTTCCAG |
| 8. pEXG2-lptD-downstream-REV | Reverse primer to amplify downstream region of native promoter of *lptD* operon. | tacaagaaagctgggtTCGCAGATCCGCTGCCAGC |
| 9. pBAD-FOR | Forward primer to amplify $P_{araBAD}$ promoter from the pUC18-derived mini-Tn7 integration vector, encoding *pmrB*, described previously [6]. | ACGCCGTTGGATACACCAAG |
| 10. pBAD-REV | Reverse primer to amplify $P_{araBAD}$ promoter from the pUC18-derived mini-Tn7 integration vector, encoding *pmrB*, described previously [6]. | GGGTATGTATATCTCCTTCTTAAAGTTAAAC |
| 11. pEXG2-lptE-upstream-FOR | Forward primer to amplify upstream region of *lptE* with pEXG2-AMP overhang for assembly of *pEXG2-AmpR-lptE* | TGCGGTATTTCACACCGCATATGCAGGAAACAGCTATGACCCGAGAAGATGTCGAAGTCG |
| 12. pEXG2-lptE-upstream-REV | Reverse primer to amplify upstream region of *lptE* with kanamycin resistance cassette overhang for assembly of *pEXG2-AmpR-lptE* | CGGCCGGAGAACCTGCGTGCAATCCATCTTGTTCAATCATCAAGCTTCCCTCACTCCTG |
| 13. pEXG2-lptE-downstream-FOR | Forward primer to amplify downstream region of *lptE* with kanamycin resistance cassette overhang for assembly of *pEXG2-AmpR-lptE* | GCGCATCGCCTTCTATCGCCTTCTTGACGAGTTCTTCTGATGCCCCGGGGCCGGTTT |
| 14. pEXG2-lptE-downstream-REV | Reverse primer to amplify downstream region of *lptE* with pEXG2-AMP overhang for assembly of *pEXG2-AmpR-lptE* | AGCATAAATGTAAAGCAAGCTTTGTAAAACGACGGCCAGTTCTCGCGGGCCAGGCCC |
| 15. pEXG2-oprL-upstream-FOR | Forward primer to amplify upstream region of *oprL* with pEXG2-AMP overhang for assembly of *pEXG2-AmpR-oprL* | TGCGGTATTTCACACCGCATATGCAGGAAACAGCTATGACCGACTACGACGGCGCC |

| | | |
|---|---|---|
| 16. pEXG2-oprL-upstream-REV | Reverse primer to amplify upstream region of *oprL* with kanamycin resistance cassette overhang for assembly of *pEXG2-AmpR-oprL* | CGGAGAACCTGCGTGCAATCCATCTTGTTCAATCATCATGTAACTCCTAATGAACCCCAG |
| 17. pEXG2-oprL-downstream-FOR | Forward primer to amplify downstream region of *oprL* with kanamycin casette overhang for assembly of *pEXG2-AmpR-oprL* | ATCGCCTTCTATCGCCTTCTTGACGAGTTCTTCTGAGAAGTCGTTATGCCCAAGCACCTG |
| 18. pEXG2-oprL-downstream-REV | Reverse primer to amplify downstream region of *oprL* with pEXG2-AMP overhang for assembly of *pEXG2-AmpR-oprL* | AGCATAAATGTAAAGCAAGCTTTGTAAAACGACGGCCAGTTGGGCGGCCGAAGTGCC |
| 19. pEXG2-FOR | Forward primer to amplify pEXG2-AMP backbone for assembly of *pEXG2-AmpR* plasmids | ACTGGCCGTCGTTTTACAAAGC |
| 20. pEXG2-REV | Reverse primer to amplify pEXG2-AMP backbone for assembly of *pEXG2-AmpR* plasmids | GTCATAGCTGTTTCCTGCATATGCG |
| 21. KanR-FOR | Forward primer to amplify KanR cassette for assembly of *pEXG2-AmpR* plasmids | ATGATTGAACAAGATGGATTGCACGC |
| 22. KanR-REV | Reverse primer to amplify KanR cassette for assembly of *pEXG2-AmpR* plasmids | TCAGAAGAACTCGTCAAGAAGGCG |
| 23. MiniTn7-backbone-FOR | Forward primer to amplify *pUC18-mini-Tn7-GmR* backbone for assembly of *pUC18-mini-Tn7-GmR-lptE, pUC18-mini-Tn7-GmR-oprL and pUC18-mini-Tn7-GmR-dxr* | AGCGCGAGTCGAGGTCGACG |
| 24. MiniTn7-backbone-REV | Reverse primer to amplify *pUC18-mini-Tn7-GmR* backbone for assembly of *pUC18-mini-Tn7-GmR-lptE, pUC18-mini-Tn7-GmR-oprL and pUC18-mini-Tn7-GmR-dxr* | GGGTATGTATATCTCCTTCTTAAAGTTAAAC |
| 25. LptE-Tn7-FOR | Forward primer to amplify *lptE* with $P_{araBAD}$ overhang for assembly of *pUC18-mini-Tn7-GmR-lptE* | GAAATAATTTTGTTTAACTTTAAGAAGGAGATATACATACCCATGAAACGTATCCTGACCAGC |
| 26. LptE-Tn7-REV | Reverse primer to amplify *lptE* with *pUC18-mini-Tn7-GmR* overhang for assembly of *pUC18-mini-Tn7-GmR-lptE* | TTAAGCTAGCTTATCGATACCGTCGACCTCGACTCGCGCTTCACGGGGTGGGGAACTC |
| 27. OprL-Tn7-FOR | Forward primer to amplify *oprL* with $P_{araBAD}$ overhang for assembly of *pUC18-mini-Tn7-GmR-oprL* | TAACTTTAAGAAGGAGATATACATACCCATGGAAATGCTGAAATTCGG |
| 28. OprL-Tn7-REV | Reverse primer to amplify *oprL* with *pUC18-mini-Tn7-GmR* overhang for assembly of *pUC18-mini-Tn7-GmR-oprL* | CGTCGACCTCGACTCGCGCTCAAGCTTATTTTACTTCTTCAGCTCGACG |
| 29. Dxr-Tn7-FOR | Forward primer to amplify *dxr* with $P_{araBAD}$ overhang for assembly of *pUC18-mini-Tn7-GmR-dxr* | TGTTTAACTTTAAGAAGGAGATATACATACCCATGAGTCGACCGCAGCG |
| 30. Dxr-Tn7-REV | Reverse primer to amplify *dxr* with *pUC18-mini-Tn7-GmR* overhang for assembly of *pUC18-mini-Tn7-GmR-dxr* | TTAAGCTAGCTTATCGATACCGTCGACCTCGACTCGCGCTCTAGCCGGCGTGTCGG |
| 31. CRISPRi-dCas9-FOR | Forward primer to amplify dCas9 with mini-Tn7 overhang for assembly of *pUC18-mini-Tn7-GmR-CRISPRI* plasmids | TTTAACTTTAAGAAGGAGATATACATACCCATGACCAACGGCAAGAT |
| 32. CRISPRi-dCas9-REV | Reverse primer to amplify dCas9 with c-terminal his6 tag for assembly of *pUC18-mini-Tn7-GmR-CRISPRI* plasmids | AATGGTGATGGTGATGATGTCCGCTTCCTGCTGCGCTTCCCTTCTTGTTGTTCTTGAAGT |
| 33. CRISPRi-miniTn7-FOR | Forward primer to amplify *pUC18-mini-Tn7-GmR backbone* for assembly of *pUC18-mini-Tn7-GmR-CRISPRi* plasmids | CAGGAAGCGGACATCATCACCATCACCATTGAAATAAGCTTTTCTAGAGCACAGC |
